# IP_3_R-TRPM4 Coupling Determines the Spatial Reach of Pericyte-Mediated Capillary Constriction

**DOI:** 10.64898/2026.06.24.734288

**Authors:** Vidya Murthy, Alex Aupetit, Ahmed Eltanahy, Albert L. Gonzales

## Abstract

Ensheathing pericytes extend multiple projections that wrap around capillary vessels to regulate diameter and direct blood flow distribution across the microvascular network. At capillary bifurcations, individual projections wrap around branches of different diameters, positioning the pericyte to simultaneously regulate multiple vessel segments. Using optogenetic tools in *acta2*-opto-α1AR and *acta2*-CatCh mice, we show that Gq-coupled receptor activation confines contractile responses to the projection receiving the stimulus, whereas direct membrane depolarization propagates to projections wrapping neighboring capillary branches via gap junction-independent mechanisms. Computational modeling revealed that TRPM4, a Ca^2+^-activated nonselective cation channel, uniquely couples local Ca^2+^ signals to voltage-gated Ca^2+^ channel activation in projections on neighboring branches at physiologically relevant channel abundances. Ca^2+^ imaging identified two kinetically distinct event populations: slow, low-amplitude IP_3_R-mediated events and fast, high-amplitude VGCC-mediated transients. Sustained low-amplitude signals selectively maintained TRPM4 activation and enabled cross-projection propagation, whereas brief high-amplitude transients drove rapid channel inactivation. Proximity ligation assays confirmed nanoscale colocalization of TRPM4 and IP_3_ receptors in pericytes. Focal IP_3_ photolysis produced constriction across projections wrapping multiple capillary branches that was abolished by TRPM4 blockade and enhanced by PKC-mediated augmentation of TRPM4 expression. These findings identify an IP_3_R-TRPM4 signaling axis as a molecular switch that gates whether capillary constriction remains branch-specific or coordinates across the pericyte, enabling precise stimulus-dependent control of blood flow distribution in the microvasculature.

## Introduction

Pericytes are specialized mural cells with octopoid-like projections that enwrap capillaries to regulate blood flow within the microvasculature. At the post-arteriolar region^1^, or capillary entry point, ensheathing pericytes^2, 3^ extend densely packed projections that completely wrap small vessel segments, with nearly half of all pericytes residing at capillary bifurcations^2, 4, 5^. This strategic positioning enables them to regulate vessel diameter and direct red blood cell distribution entering the capillary network^5^. Each projection functions as an independent signaling unit capable of rapid, localized Ca^2+^ dynamics and contractility^5^, a property that fundamentally distinguishes pericytes from vascular smooth muscle cells (vSMCs), which contract as a single circumferential unit. This distinction has important mechanical consequences. According to Laplace’s law (T = ΔP·R), circumferential wall tension is proportional to both transmural pressure and vessel radius, meaning that mural cells on larger vessels experience greater stretch than those on smaller ones. In vSMCs, this tension is distributed uniformly along the cell’s long axis, producing a globally integrated mechanical response. In ensheathing pericytes, by contrast, each projection independently wraps around the vessel wall and senses wall tension in isolation, a difference that becomes even more pronounced in junctional pericytes, whose projections span capillary branches of unequal diameter and are therefore simultaneously exposed to different levels of circumferential stretch. Despite this capacity for local autonomy, pericyte contraction^6^ ^7^, like that of SMCs^8^ ^9, 10^, is ultimately triggered by Ca^2+^ influx through voltage-gated Ca^2+^ channels (VGCCs)^11^. This raises a fundamental question: how do ion channels within individual pericyte projections maintain the signaling autonomy needed to sense and respond to local capillary branch activity while still coordinating with neighboring projections to generate integrated vascular responses?

Ion channels rarely function in isolation. In excitable and contractile cells, they operate in series or in parallel, with the output of one channel serving as the input for another to orchestrate complex responses across space and time. In vSMCs, transient receptor potential (TRP) channels exemplify this principle: TRPC3 and TRPC6 act as receptor- and pressure-operated channels that mediate localized Ca^2+^ entry^12, 13^, while TRPM4, though impermeable to Ca^2+^, translates these local Ca^2+^ elevations into membrane depolarization via Na^+^ influx^14, 15^, which in turn activates VGCCs to amplify and synchronize Ca^2+^ influx and drive contraction^16^. Together, these channels form interconnected signaling circuits that integrate receptor activation, mechanical stimuli, membrane excitability, and Ca^2+^ dynamics to coordinate vascular tone. Whether analogous ion channel circuits operate in pericytes, and if so, how they are organized across morphologically distinct projections to simultaneously support local contractile autonomy and inter-projection coordination, remains unknown. Here, we address this question using a combination of optogenetic and pharmacological tools, Ca^2+^ imaging, local IP_3_ photolysis, and computational modeling to dissect the principles governing subcellular signal propagation in ensheathing pericytes. We demonstrate that signals initiated in one projection can propagate to neighboring projections through a TRPM4-dependent mechanism, and that the spatial reach of this propagation is dynamically gated by the nature and context of the initiating stimulus. Our findings reveal that ion channels in pericyte projections do not operate independently but rather as part of a spatially organized signaling architecture, operating in series, in parallel, and in functional opposition, to enable precise, branch-specific regulation of capillary perfusion.

## METHODS

### Animals

All animal studies were performed in accordance with the guidelines of the Institutional Animal Care and Use Committee (IACUC) under Protocol No. 20-07-1039-1 at the University of Nevada, Reno and Protocol No. 17-034 at the University of Vermont. Adult (2–5-month-old) male and female mice were group-housed with environmental enrichment and free access to food and water. The following mice strains were used C57BL/6J (Stock No. 000604; Jackson Laboratories); *NG2*-dsRed (Stock No. 008241; Jackson Laboratories); acta2-opto-α1AR (acta2-opto-α1AR-IRES-lacZ)^17^; acta2- CatCh (acta2-CatCh-IRES-lacZ)^17^; and acta2-GCaMP-mCherry^17^ mice were originally obtained from the CHROMus resource and are available in Jackson Laboratories (Stock No. 028346, 028348, and 025405 respectfully). Animals were euthanized either by deep anesthesia with pentobarbital sodium (50 mg, i.p.) followed by exsanguination and decapitation, or by 4% isoflurane anesthesia followed by rapid decapitation. Retinas were then isolated, mounted in an *en face* configuration (vitreal side facing upward), pinned in place, and maintained in ice-cold retinal physiological saline solution.

### Solutions and Reagents

Retina dissection buffer consists of 119 mM NaCl, 3 mM KCl, 0 mM CaCl_2_, 3 mM MgCl_2_, 5 mM glucose, 26.2 mM NaHCO_3_ and 1 mM NaH_2_PO_4_, bubbled with 95% O_2_/5% CO_2_ (pH 7.4). The tissue was maintained in a saline solution with HEPES buffer to maintain physiological pH (124 mM NaCl, 10 mM HEPES, 1mM NaH_2_PO_4_, 2.5 mM KCl, 1.8 mM CaCl_2_, 2 mM MgCl_2_, and 10 mM glucose). All chemicals and drugs were purchased from Sigma-Aldrich (St. Louis, MO, USA) unless specified otherwise.

### *Ex vivo* Retinal Preparations

*En face* retinal preparations were generated as previously described^5, 7^. Briefly, following euthanasia, eyes were enucleated and transferred to ice-cold physiological saline solution (PSS). The cornea, lens, and vitreous humor were removed, and the retina was dissected free from the retinal pigment epithelium and sclera. Retinas were mounted *en face* with the vitreal surface facing upward on a silicone block and maintained in ice-cold Mg-PSS (5 mM KCl, 140 mM NaCl, 2 mM MgCl_2_, 10 mM HEPES, 10 mM glucose).

For optogenetic stimulation, whole-mount retinas from acta2-opto-α1AR and acta2-CatCh mice were pinned *en face* in a superfused recording chamber and perfused with oxygenated rPSS at 36°C. Capillary networks were visualized by fluorescence microscopy with isolectin B4 labeling. Simultaneous imaging and focal photostimulation were performed using a Yokogawa spinning-disk confocal (Andor Revolution) with a Mosaic digital illumination system on an upright Nikon microscope (60×, NA 1.0), with 488/560 nm excitation and appropriate band-pass emission filters. Focal 405 nm illumination activated the opto-α1AR fusion protein (DAG/IP_3_ signaling) or CatCh (membrane depolarization) in targeted pericyte projections. Capillary diameters were measured before, during, and after stimulation.

For Ca^2+^ imaging, as previously described^5, 7^, *en face* retinas from *acta2*-GCaMP-mCherry mice were prepared and perfused as above, then stained with rhodamine- or FITC-conjugated isolectin B4 (1:25, 20 min, 37°C). Imaging was performed on the same spinning-disk confocal system (60x, NA 1.0) at 512 × 512 pixels, 30 frames/s. GCaMP reported intracellular Ca^2+^ dynamics, while mCherry served as a motion artifact control. Ca^2+^ transients were quantified as F/F_0_, with event duration reported as t_1/2_ and frequency as events per recording interval, analyzed using SparkAn (Nelson Lab, Burlington VT) and ImageJ (1.54, NIH). To characterize the kinetic properties of Ca^2+^ events in ensheathing pericytes, recordings obtained from these preparations during pharmacological experiments were re-analyzed from the dataset reported in Gonzales et al. (2020)^5^. Individual Ca^2+^ events were identified from pericyte projections under baseline and pharmacological conditions, and peak fluorescence intensity (F/F_0_) was plotted against half-peak duration (t_1/2_) to characterize the amplitude and kinetic profile of distinct Ca^2+^ event populations, a question not addressed in the original publication.

For pressurized-retina studies, as previously described^5, 7^, the intact orbit with optic nerve, ophthalmic artery, and musculature was isolated in ice-cold, Ca^2+^-free, Mg^2+^-supplemented dissection solution (124 mM NaCl, 26 mM NaHCO_3_, 1 mM NaH_2_PO_4_, 2.5 mM KCl, 3.8 mM MgCl_2_, 10 mM glucose). After removing surrounding tissues, off-target ophthalmic artery branches were ligated, and the artery was cannulated with a glass cannula attached to a micromanipulator. The retina was flattened on a custom silicone platform and superfused with oxygenated rPSS at 37°C (5 mL/min). Intraluminal pressure was controlled via a gravity-fed system and pressure monitor (PM-4; Living Systems Instrumentation), with pressurization to ∼60 mmHg confirmed by clearance of blood cells from the retinal vasculature.

For IP_3_ photo-uncaging experiments, wildtype C57BL/6J retinas were imaged on a Crest Optics X-Light V3 spinning-disk confocal equipped with a 7-wavelength laser launch (LDI-7; 89 North) and dual sCMOS cameras (ORCA-Fusion Gen-III; Hamamatsu) on an Olympus BX51WI microscope. IP_3_ uncaging was achieved using a Mightex Polygon DMD illuminator (Polygon1000-G) with 405 nm illumination targeted to pericyte-containing branching zones. Imaging was performed on the same spinning-disk confocal system (60x, NA 1.00) at 512 × 512 pixels, 30 frames/s. Capillary diameters were measured before, during, and after stimulation; the stimulated branch was designated B_S_ and unstimulated branches as B_NS_.

### Pharmacology

Retinas were treated with pharmacological modulators of TRPM4-, TRPC3/6-, VGCC-, IP_3_R-, and PKC-dependent signaling pathways identified in the experimental figures. Preparations were incubated with the TRPM4 activator PMA (1–3 µM, Santa Cruz Biotechnology), TRPM4 inhibitors 9-phenanthrol (30 µM, Sigma) and NBA (1–5 µM, Glixx Laboratories), TRPC3/6 inhibitor Pyr3 (10 µM, Tocris), TRPC3 inhibitor GSK2833503A (3–100 nM, Tocris), TRPC3 agonist GSK1702934A (100 nM, Tocris), VGCC inhibitor nimodipine (100 nM, Sigma Aldrich), SERCA inhibitor cyclopiazonic acid (CPA; 10–50 µM, Tocris), IP_3_ receptor inhibitor xestospongin C (1 µM, Cayman Chemical) and ANO1 inhibitor Ani9 (5 µM, Cayman Chemical). Additional treatments included the thromboxane A_2_ receptor agonist U46619 (100 nM, Cayman), membrane- permeable Bt-IP_3_ (10 µM, SiChem), extracellular KCl (60 mM), and extracellular Ca^2+^-free solution. For IP_3_ uncaging experiments, retinas were incubated with caged IP_3_ (ci-IP_3_/PM; 1 µM, Tocris, 30 min). Isolectin B4 (Griffonia simplicifolia lectin I, Vector Laboratories) was included during incubations to visualize the retinal vasculature. All treatments were performed at 37°C unless otherwise indicated.

### Computation Modeling

The model presented here is a compartmentalized two projection pericyte model which are electrically coupled together (Projection 1, P1 and Projection 2, P2). When one projection is activated by intracellular Ca^2+^ signal followed by IP_3_ receptor mediated Ca^2+^ release, the signal propagates downstream to P2 and the subsequent entry of Ca^2+^ to P2 through VGCCs. Each projection is modeled as an individual compartment capable of contracting independent of neighboring projections within the same cell. The projections are connected through axial conductance and membrane potentials. Simulations are implemented in Python using custom algorithms. NumPy^18^ (v2.0.2, NumFOCUS, Austin, TX, USA) and Matplotlib^19^ (v3.9.4, Matplotlib Development Team, numfocus.org) were used for mechanistic calculations and visual graphing. Time integrations were performed using Euler method^20^ with a time step of 0.1 ms. The unit for the voltages is in millivolts (mV), Ca^2+^ concentrations in micromolar (uM), conductance in picosiemens (pS) and resistance in Ohms (W). Intracellular Ca^2+^ concentration is modeled to be a dynamic entity, which allows for understanding of how Ca^2+^ signal waveform regulates TRPM4 activation. To maintain physiological consistency, we capped Ca^2+^ concentration at a maximum of 1 uM to maintain physiologically relevant range. TRPM4 is modeled as a Ca^2+^ activate, non-selective cation channel with maximum conductance at 30 pS^14, 21^. TRPC3 is modeled as a DAG-activated non-selective cation channel with maximum conductance at 60 pS^12, 22^. ANO1 is modeled as Ca^2+^ activated Cl^-^ channel with maximum conductance at 2 pS^23–25^. Membrane potential evolves in accordance with TRPM, TRPC3 or ANO1 mediated currents depending on the simulation, VGCC activation in P2 is modeled as a voltage dependent conductance controlled by membrane potential. To examine how channel expression influences downstream signaling, simulations were repeated with wide range of TRPM4, TRPC3 or ANO1 densities, the outputs were quantified and plotted.

### Immunocytochemistry and Proximity Ligation Assay (PLA)

For immunocytochemistry experiments, retinas were processed as previously described^5^. Briefly, retinas were fixed in 4% paraformaldehyde prepared in Mg-PSS for 15 min, permeabilized with 0.1% Triton X-100, and blocked in Mg-PSS containing 2% bovine serum albumin and 2% normal goat serum. Retinas were incubated overnight at 4°C with primary antibodies diluted in blocking solution containing goat anti-TRPM4 (1:50; Santa Cruz Biotechnology, sc-27540) and anti-α-smooth muscle actin-Alexa Fluor 420 (1:50; Invitrogen, 53-9760-82). Preparations were washed in Mg-PSS and incubated for 2 h at room temperature with FITC-conjugated donkey anti-goat secondary antibody (1:200; Santa Cruz Biotechnology, sc-2024). Retinas were subsequently washed, mounted using Fluoromount-G, and imaged using spinning-disk confocal microscopy.

For PLA experiments, following harvest, retinas were fixed in 4% paraformaldehyde for 20 minutes. Neuronal tissue digestion was carried out by incubating retinas in ddH_2_O for 1 hour, followed by treatment with TrypLE™ Express Enzyme (ThermoFisher, USA) for 1.5 hours at 37°C. Bruch’s membrane was removed, and the vascular tissue was gently triturated by pipetting to eliminate residual neuronal components. The isolated vasculature was transferred onto a glass slide and air-dried. Proximity ligation assay (PLA) was performed using the Naveni TriFlex Cell kit according to the manufacturer’s protocol. Two primary antibody pairs were used: TRPM4 (Alomone Labs, USA)–IP_3_R (Abcam, USA) and TRPM4–Ca_V_1.2 (NeuroMab, California, USA). Imaging was conducted using a Leica Stellaris 8 confocal microscope, and image analysis was performed with Imaris (Oxford Instruments, USA) and Fiji. Images were processed and analyzed using ImageJ. For proximity ligation assays (PLA), a vascular mask was generated from the lectin image. Regions of interest (ROIs) corresponding to arteries and capillaries were defined, and object detection was applied to identify, count, and measure PLA puncta.

### Statistics

Data in figures and text are presented as means ± standard error of the mean (SEM). Values of “n” refer to the number of experiments, and “N” refers to the number of animals. Statistical analyses were performed using Prism 10 (GraphPad). Statistical significance was determined using unpaired student’s t-tests, one-way analysis of variance (ANOVA), two-way repeated-measures ANOVA. P-values ≤ 0.05 were considered statistically significant for all experiments.

## RESULTS

### Ensheathing pericyte projections exhibit localized GPCR signaling, yet neighboring projections are coupled through membrane depolarization

Ensheathing pericytes consist of a cell body that sits atop the capillary vessels with extensions along the vessels and projections that wraparound the capillary vessels. A majority of pericytes are positioned at capillary junctions^2, 4, 5^, allowing their projections to wrap around multiple capillary segments (Fig. 1A). Previous work from our lab demonstrated the capacity of individual pericyte projections to compartmentalize their Ca^2+^ and contractile dynamics contributing to branch-specific constriction of different capillary branches^5^. To examine how signals propagate between pericyte projections, we used two optogenetic mouse models, *acta2*-opto-α1AR and *acta2*-CatCh, in which the *acta2* (α actin) promoter drives expression in smooth muscle cells and ensheathing capillary pericytes^17^. In these models, blue light either activates Gq coupled α1 adrenergic receptors (opto-α1AR) or directly depolarizes the membrane via a light gated cation channel (CatCh). Activation of Gq-coupled G protein-coupled receptors (GPCRs) stimulates phospholipase C, leading to the local generation of inositol 1,4,5-trisphosphate (IP_3_) and diacylglycerol (DAG); IP_3_ mobilizes Ca^2+^ release from intracellular stores, while DAG activates protein kinase C (PKC), together initiating downstream signaling cascades that regulate excitability and contraction. For this experimental series, we identified a single junctional pericyte (Fig. 1A; schematical representation indicates the pericyte nucleus) based on two criteria: first, neighboring pericyte nuclei were located more than 10 µm away; and second, the pericyte extended projections that enwrapped different capillary branches, as determined from z-stack imaging. Using an inline pattern illuminator, which provides precise spatiotemporal and high-resolution control of light delivery, we photostimulated a region of interest (ROI) corresponding to a single branch of the capillary junction and measured diameter changes in the two adjoining branches. Using this approach, we selectively induce local DAG and IP_3_ release within individual projections and constriction of the stimulated capillary branch (B_S_), then assess whether these signals propagate to projections of the same pericyte on other non-stimulated capillary branches (B_NS_), resulting in constriction of those vessel segments. In retinas isolated from *acta2*-opto-α1AR mice, 405 nm stimulation induced constriction of the stimulated branch (measured as percent constriction), while no significant changes in capillary diameter were observed in the non-stimulated branches (Fig. 1B-C). To confirm that this constriction was mediated by α1AR signaling, we bath-applied prazosin, a selective α1AR antagonist^26^, which abolished the light-evoked constriction of the stimulated branch (Fig. 1D). To assess the contribution of intracellular Ca^2+^ store release, we applied cyclopiazonic acid (CPA), a potent SERCA pump inhibitor that depletes ER Ca^2+^ stores^27^, had no effect on the constriction response. Removal of extracellular Ca^2+^ ([Ca^2+^]_o_=0) similarly eliminated the constriction response, confirming a Ca^2+^ influx-dependent mechanism. Pharmacological inhibition of TRPM4 (9-phenanthrol^27^) or L-type VGCCs (nimodipine^28^) did not significantly affect light-evoked constriction of the stimulated branch; however, selective blockade of TRPC3 with PYR3^29^ significantly attenuated this response (Fig. 1D). Together, these findings indicate that DAG and IP_3_ signaling, acting locally through TRPC3, remains confined to the stimulated projection and does not propagate to neighboring projections.

**Fig. 1.**
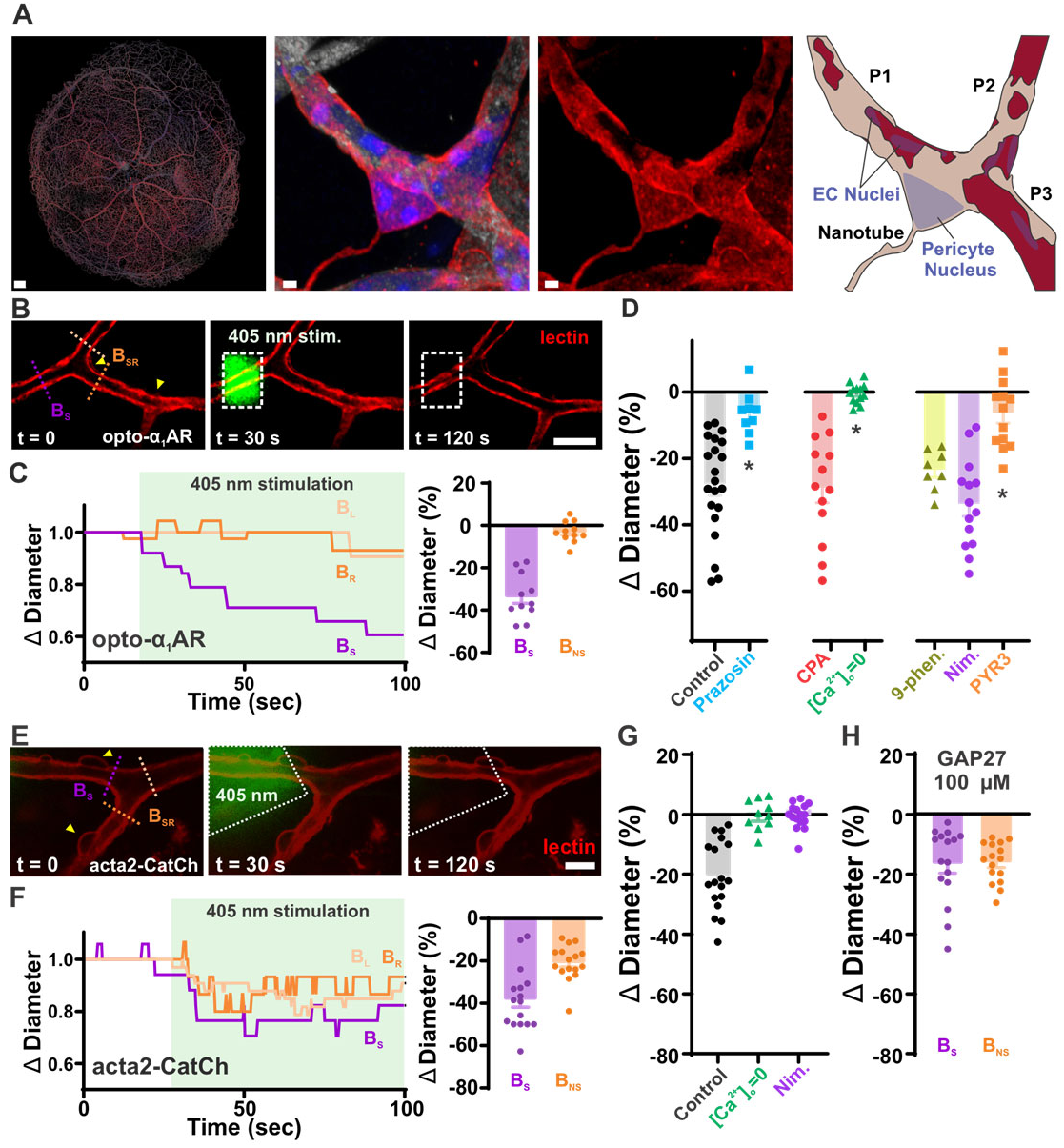
Optogenetic dissection of Gq-coupled receptor and membrane depolarization signaling reveals distinct spatial reach of pericyte contractile responses in the retinal microvasculature. **A, left**: Representative fluorescence image of a all retinal vasculature (CD31, grey) and pericyte coverage (NG2, red) capture by confocal microscopy. Scale bar, 200µm. **A, center**: Zoom on an ensheathing pericyte, stained with NG2 (red), illustrating the morphological organization of projections wrapping individual capillary, stained with CD31 (grey) branches at a bifurcation point. Nuclei are stained with DAPI (blue). The first image is a merge of all chanels and the second image is NG2 isolated to highlight pericyte morphology. Scale bar, 5µm. **A, right**: Schematical view of the pericyte (tan) wrapping around the vasculature (red) the pericyte projections are noted P1, P2 and P3. **B:** Representative confocal images of a retinal whole-mount preparation from an *acta2*-opto-α1AR mouse before (t = 0), during (t = 30 s), and after (t = 120 s) focal 405 nm photostimulation (dashed box). The stimulated branch (B_S_) and non-stimulated branches (B_SR_) are indicated. Lectin labeling (red) delineates the capillary lumen. Scale bar, 10 µm. **C, left:** Representative time course of normalized capillary diameter in stimulated (B_S_, purple) and non-stimulated branches (B_NS_, orange) during 405 nm stimulation (shaded region) in *acta2*-opto-α1AR retinas. **C, right:** Summary data of mean diameter changes in stimulated (B_S_) and non-stimulated (B_NS_) branches (right). **D:** Summary data showing pharmacological dissection of light-evoked constriction in *acta2*-opto-α1AR retinas. Prazosin (100 nM, α1AR antagonist), cyclopiazonic acid (CPA;10 µM, SERCA pump inhibitor), and removal of extracellular Ca^2+^ ([Ca^2+^]_o_=0) each abolished constriction. Selective TRPC3 blockade with PYR3 (10 µM) significantly attenuated constriction, whereas inhibition of TRPM4 (9-phenanthrol, 30 µM) or L-type VGCCs (nimodipine, 100 nM) had no significant effect. n = 8-21 pericytes per group. **E:** Representative confocal images of a retinal whole-mount from an *acta2*-CatCh mouse before (t = 0), during (t = 30 s), and after (t = 120 s) focal 405 nm photostimulation. Scale bar, 10 µm. **F, left**: Representative time course of normalized capillary diameter in B_S_ (purple) and B_SR_ (orange) branches during 405 nm stimulation in *acta2*-CatCh retinas. **F, right**: (left). Summary data of mean diameter changes in B_S_ and B_NS_ branches (right). **G:** Summary data showing pharmacological dissection of CatCh-evoked constriction. Removal of extracellular Ca^2+^ abolished stimulated branch constriction, while nimodipine attenuated constriction in both stimulated and non-stimulated branches, implicating VGCC activation in the propagated response. n = 10-18 pericytes per group. **H:** Summary data showing the effect of GAP27 (100 µM), a connexin mimetic peptide blocking gap junction communication, on CatCh-evoked constriction. GAP27 had no significant effect on constriction in either branch, indicating that depolarization propagation between pericyte projections does not require endothelial gap junction coupling. n = 12 pericytes. Data are presented as means ± SEM. Paired t-tests were used for comparisons between stimulated and non-stimulated branches (C, F, and H); one-way ANOVA used for pharmacological comparisons against control (D and G). *P ≤ 0.05 vs control. N = 3-5 mice per experiment.

We next investigated whether membrane depolarization could propagate between branches using the *acta2*-CatCh mouse model, which expresses a modified channelrhodopsin-2 (ChR2), a blue light–activated non-selective cation channel (activation spectrum 400–520 nm)^30^. Light-evoked cation entry depolarizes the membrane, activating VGCCs and driving contraction. Following photostimulation, we observed constriction of both the stimulated branch (B_S_: ∼−35–40%) and non-stimulated branches (B_NS_: ∼−20–25%) of comparable magnitude (Fig. 1E–F). As observed with opto-α1AR stimulation, removal of extracellular Ca^2+^ abolished constriction of the stimulated branch, while nimodipine also significantly attenuated constriction, consistent with VGCC-mediated Ca^2+^ influx driving the contractile response (Fig. 1G). To determine whether ChR2-mediated membrane depolarization propagates to non-stimulated branches via the underlying endothelial cell network, we applied GAP27, a connexin mimetic peptide that blocks gap junction communication^31^. GAP27 (100 µM) had no effect on constriction in either stimulated or non-stimulated branches, suggesting that depolarization does not spread through the endothelium, but rather propagates directly between pericyte projections (Fig. 1H). Together, these results demonstrate that pericyte contractile signaling is modality-dependent: local GPCR activation drives TRPC3-dependent constriction confined to the stimulated projection, whereas membrane depolarization propagates directly between pericyte projections independent of the endothelium, activating VGCCs and coordinating constriction across multiple capillary branches of a single pericyte.

### Ensheathing Pericytes Express TRPM4, and Modeling Reveals Its Role in Projection-to-Projection VGCC-activation

Because membrane depolarization, but not second messenger signaling, can rapidly spread between pericyte projections, we tested whether localized activation of GPCR-linked ion channels can trigger voltage-gated Ca^2+^ channel (VGCC) activation in a neighboring projection. We developed a simplified two-compartment model of electrically coupled pericyte projections. Depolarizing ion channels (TRPM4, TRPC3, or ANO1; tested separately) were activated in Projection 1, and passive voltage spread to Projection 2 was used to assess VGCC activation. Membrane properties^14,32, 33, 34^, VGCC density^8, 11, 35, 36^, and stimulus amplitude were held constant to isolate each channel’s depolarizing contribution. VGCC activation was modeled using established methods^20, 32, 33, 37–39^ with Boltzmann kinetics consistent with L-type Ca^2+^ channels (Ca_V_1.2)^35^. Signal propagation was quantified by VGCC activation in Projection 2 under physiologically relevant conditions^2, 30, 39–41^, similar to compartmental models of small cellular processes^39, 42^. Ion channel parameters, including conductance, reversal potential, resting membrane potential, and axial resistance, were derived from experimental data^12, 21, 23, 39, 43–45^ ^3^ ^39, 46–48^. TRPM4 was modeled as a Ca^2+^-activated monovalent cation channel (∼20–30 pS, ∼0 mV reversal)^14, 37^. TRPC3 as a DAG-activated non-selective cation channel (∼40–60 pS)^12, 22^, and ANO1 as a Ca^2+^-activated Cl^-^ channel (∼1–3 pS, ∼−30 mV reversal)^23–25^. Qualitative differences between channel types were independent of resistance values.

Using this model, we observed that trans-projection activation of VGCC depended strongly on channel type and abundance. Increasing TRPM4 number produced a steep rise in VGCC conductance, reaching near-maximal activation with relatively few channels before rapidly saturating (Fig. 2A). In contrast, TRPC3, a non-selective cation channel expressed in pericytes^45, 49^, produced a modest, gradual increase in VGCC conductance even at high channel numbers (Fig. 2A; Sup. Figure 1); at extreme TRPC3 densities (100–2000 channels), VGCC conductance only approached saturation, confirming that TRPC3 requires channel densities orders of magnitude greater than TRPM4 to achieve equivalent downstream activation (Sup. Figure 2), while ANO1, a Ca^2+^-activated Cl^-^ channel also expressed in pericytes^25, 45^, had minimal effect across the tested range. These results reveal marked differences in the efficiency with which individual ion channels couple local Ca^2+^ signals to propagated depolarization. TRPM4 was uniquely effective, generating sufficient depolarization to robustly activate VGCCs in a neighboring projection with far fewer channels than TRPC3 and substantially outperforming ANO1. Given this pronounced efficiency, we prioritized TRPM4 for further experimental and computational investigation as a key mediator of electrical communication between pericyte projections.

**Figure 2:**
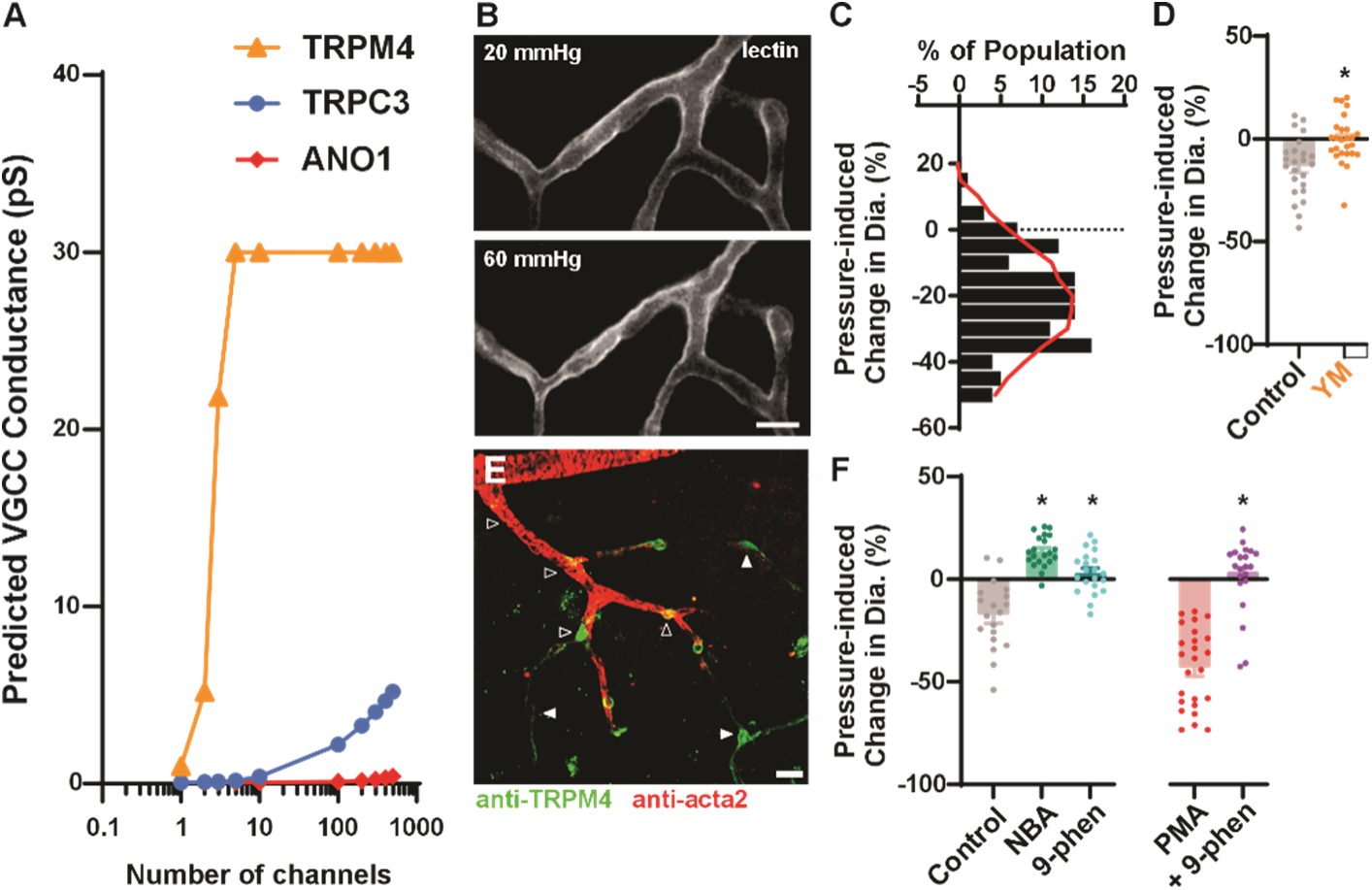
TRPM4 is expressed in ensheathing pericytes of the retinal microvasculature. **A:** Computational modeling predicting VGCC conductance in a distal pericyte projection (P2) as a function of channel number for TRPM4 (orange), TRPC3 (blue), and ANO1 (red). TRPM4 achieves maximal VGCC conductance (∼30 pS) at substantially lower channel abundance than TRPC3 or ANO1, providing a mechanistic rationale for TRPM4 as the primary candidate for inter-projection electrical coupling. **B:** Representative confocal images of a pressurized *ex vivo* retinal preparation at 20 mmHg (top) and 60 mmHg (bottom) with lectin labeling (gray) delineating the capillary lumen. Elevation of intraluminal pressure to 60 mmHg produced a measurable reduction in capillary diameter. Scale bar, 10 µm. **C:** Frequency distribution of pressure-induced diameter changes (%) across the capillary population (n = 111 measurements), with the distribution curve (red) skewed toward constriction, confirming that ensheathing pericytes constrict in response to elevated intraluminal pressure. **D:** Summary data of pressure-induced diameter changes in control and YM-254890 (YM; 1 µM)-treated preparations, implicating GqPCR-dependent signaling in the pericyte pressure-sensing response. n = 24-26 pericytes per group. **E:** Representative confocal image of a retinal whole-mount immunolabeled for TRPM4 (green) and acta2 (red), demonstrating TRPM4 protein expression in ensheathing pericytes (open arrowheads) and thin-stranded pericytes (closed arrowheads) along the retinal capillary network. Scale bar, 10 µm. **F:** Summary data showing that pharmacological inhibition of TRPM4 with NBA (1 µM) or 9-phenanthrol (30 µM) significantly attenuated pressure-induced constriction (top), while PKC activation with PMA (1 µM) enhanced constriction in a TRPM4-dependent manner, as evidenced by abolition of the PMA effect by co-application of 9-phenanthrol (30 µM) (bottom). n = 19-21 pericytes per group. All data are presented as means ± SEM. Comparisons between two groups were made using unpaired t-tests; multiple group comparisons were made using one-way ANOVA. P ≤ 0.05 versus control. N = 3-5 mice per condition.

In vascular SMCs, increases in arterial pressure can activate multiple GqPCRs, leading to elevations in intracellular Ca^2+^ and activation of depolarizing ion channels, including TRPM4, which promote membrane depolarization and vasoconstriction^13, 50^. To examine the effect of intraluminal pressure on the contractile state of ensheathing pericytes, we performed a series of experiments using our pressurized-retina preparation^7^. Isolated retinas were pinned down *en face* and the feeding ophthalmic artery was cannulated and pressurized. Consistent with previous findings^6, 7^, increasing intraluminal pressure from 20 to 60 mmHg induced constriction in all capillary vessels covered by ensheathing pericytes. To quantify how much of the vessel was constricted by ensheathing pericytes and to what extent, we measured the diameter of capillary vessels at ∼5-µm increments along the capillary vessels (Fig. 2B). Increases in intraluminal pressure led to a decrease in average measured diameter, as depicted in our histogram (Fig. 2C), suggesting a uniform response throughout the capillary segment. Pre-incubation with the Gα_q/11_-selective GPCR blocker, YM-254890 (1 µM for 10-15 mins) prevented pressure-dependent vasoconstriction (Fig. 2D), suggesting a role for GqPCRs in initiating the pressure transduction pathway in ensheathing pericytes. Next, we examined the role of TRPM4 in pressure-induced constriction of capillary vessels. Using two TRPM4 inhibitors, 4-Chloro-2-(1-naphthyloxyacetamido)benzoic acid ammonium salt (NBA, 1 µM)^51^ and 9-phenanthrol^52^ blocked the pressure-induced constriction of ensheathing pericytes (Fig. 2E). As previously reported in vascular SMCs^53^, we propose that activation of protein kinase c (PKC) will increase TRPM4 activity in pericytes by increasing its surface expression. In support of this hypothesis, pressure-induced vasoconstriction was enhanced following activation of PKC with PMA (phorbol 12-myristate 13-acetate, 1 µM). This effect was blocked by 9-phenathrol (Fig. 2F), suggesting that the PMA-induced increase in vascular reactivity is attributable to increased TRPM4 channel activity. Consistent with previous studies^7, 45^, immunohistochemical staining confirmed TRPM4 channel expression in pericytes (Fig. 2E). These data support a role for TRPM4 channels in modulating the pressure transduction pathway in ensheathing pericytes.

### Modeling Ca^2+^-dependent activation of TRPM4 to electrically couple pericyte projections

In whole-cell electrophysiology, rupturing the membrane beneath the patch pipette provides access to the full intracellular space, allowing measurement of summed ionic currents across all active channels in the membrane. While such recordings have characterized ion channel activity in vSMCs^9, 11, 54^, ensheathing pericyte projections each operate as independent signaling domains. Consequently, whole-cell recordings would aggregate electrical currents from the whole cell, potentially obscuring the localized signaling events and distinct electrical activity occurring both within individual pericyte projections and between neighboring projections. To address this gap, we propose a mechanistic computational model that simulates the activation of TRPM4 from an intracellular Ca^2+^ increase, allowing us to quantify membrane depolarization and activation of VGCCs, in distant projections.

TRPM4 channels are Ca^2+^-activated ion channels, and multiple Ca^2+^-dependent mechanisms have been implicated in their physiological activation across different cell types^14, 55^. As a first step toward defining the sources and kinetics of Ca^2+^ signals in ensheathing pericytes, we re-analyzed Ca^2+^ recordings from individual pericyte projections in acta2-GCaMP-mCherry retinas originally reported in Gonzales et al. (2020)^5^. Whereas the original study characterized Ca^2+^ event frequency in response to pharmacological stimuli, here we examined the amplitude and kinetic profile of individual Ca^2+^ events to determine whether distinct event populations could be identified. From each recording, individual Ca^2+^ events were identified and peak fluorescence intensity (F/F_0_) was plotted against half-peak duration (t_1/2_) under control conditions, revealing two distinct populations of Ca^2+^ events: low-amplitude, long-duration (slow) events, and high-amplitude, short-duration (fast) events (Fig. 3B–C; Sup. Figure 4). To identify the sources of these distinct event populations, we applied pharmacological interventions targeting Ca^2+^ influx and intracellular Ca^2+^ release pathways. Application of the thromboxane A2 receptor agonist U46619 (100 nM), which activates Gq-coupled signaling and stimulates IP_3_ production, increased the frequency of slow and fast events (Fig. 3D). Membrane depolarization with 60 mM KCl increased the frequency of fast, high-amplitude events, while subsequent inhibition of L-type Ca^2+^ channels with nimodipine (100 nM) selectively eliminated these fast events, shifting the population toward slow events exclusively (Fig. 3E). To test whether IP_3_-mediated Ca^2+^ release accounted for the slow events, we applied the membrane-permeable IP_3_ analog Bt-IP_3_ (10 µM)^40, 56^, which increased the overall frequency of Ca^2+^ events, predominantly augmenting the slow, low-amplitude population (Fig. 3F). Consistent with this interpretation, inhibition of IP_3_ receptors with xestospongin C (1 µM) blocked the slow events, leaving only the fast, high-amplitude population (Fig. 3F). Together, these results indicate that pericyte Ca^2+^ dynamics involve at least two distinct signaling modes: IP_3_R-mediated release of Ca^2+^ from intracellular stores, which contributes to slow, low-amplitude events, and Ca^2+^ influx through L-type VGCCs, which contributes to fast, high-amplitude events, though the contribution of additional ion channels cannot be excluded.

**Figure 3:**
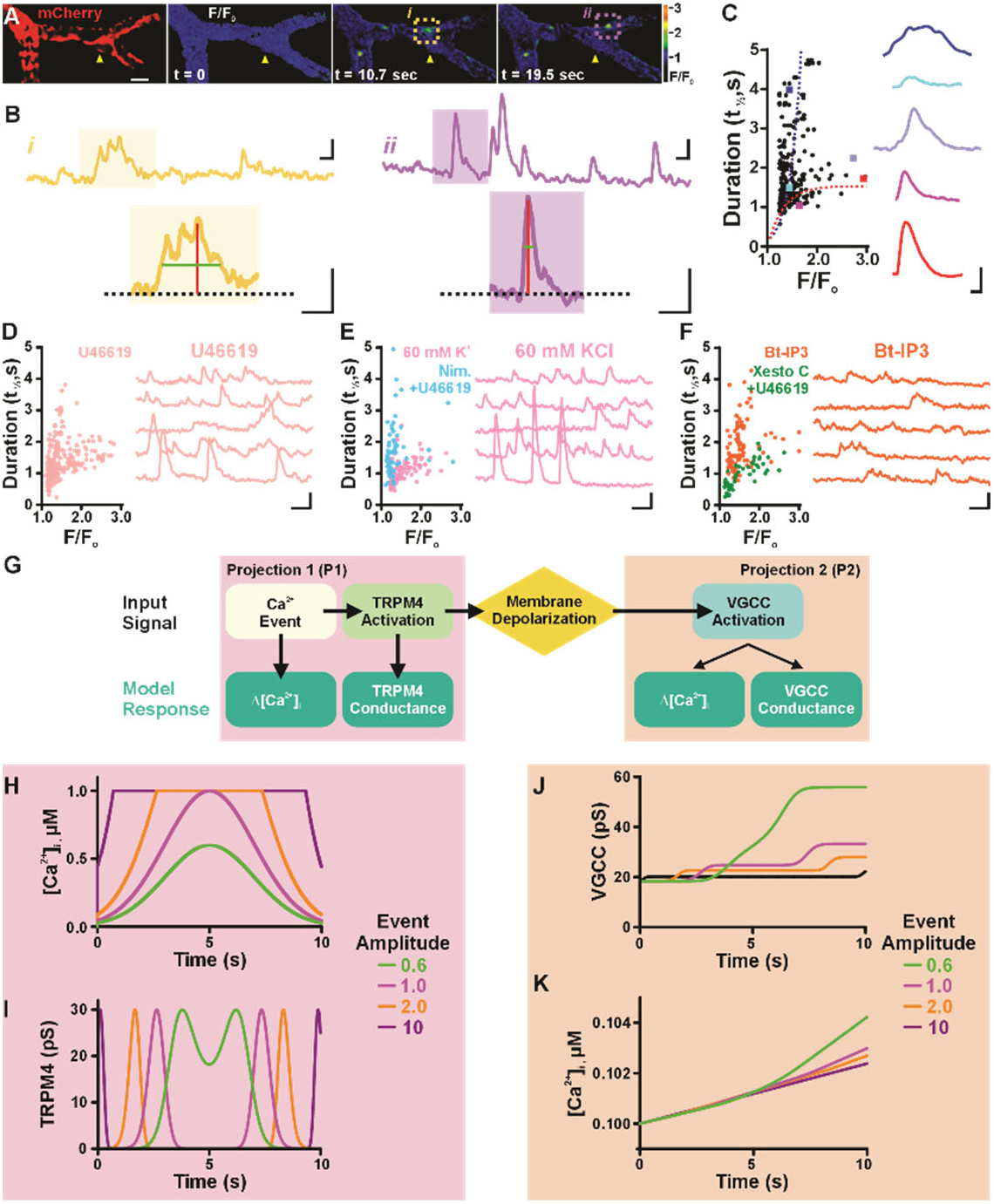
Pericyte Ca^2+^ event kinetics determine the efficacy of TRPM4-dependent inter-projection signal propagation. **A:** Representative fluorescence images of spontaneous Ca^2+^ transients recorded from individual pericyte in retinal whole-mount preparations from *acta2*-GCaMP-mCherry mice. GCaMP fluorescence (blue/purple) reports Ca^2+^ dynamics and mCherry (red) serves as a Ca^2+^-independent structural reference. Yellow arrowhead indicates the cell body. Scale bar: 10 µm. **B:** Representative Ca^2+^ fluorescence traces (F/F_0_) from individual pericyte projections under control conditions illustrating low-amplitude, long-duration (slow) events (yellow, left) and high-amplitude, short-duration (fast) events (purple, right). Insets show representative single events on an expanded timescale. Scale bars: 2 s (horizontal), 0.2 F/F_0_ (vertical). n = 327 events **C:** Scatter plot of Ca^2+^ event peak amplitude (F/F_0_) versus half-peak duration (t_1/2_) under control conditions. Representative event waveforms for each population are shown to the right, color-coded by amplitude. Scale bars: 1 s (horizontal), 1.0 F/F_0_ (vertical). **D:** Scatter plot and representative Ca^2+^ events following the exposure to the thromboxane A2 receptor agonist U46619 (100 nM). n = 341 events **E:** Scatter plot and representative Ca^2+^ events following membrane depolarization with 60 mM KCl (pink) compared to the co-application of nimodipine (Nim.; 100 nM) and U46619 (100 nM) in light blue. n = 71-101 events. **F:** Scatter plot and representative Ca^2+^ events following the application of the membrane-permeable IP_3_ analog Bt-IP3 (10 µM, orange) compared to co-application of xestospongin C (Xesto C; 1 µM) and U46619 (100 nM) in green. Scale bars (D-F): 5 s (horizontal), 0.5 F/F_0_ (vertical). n = 35-53 events. N = 3-5 mice per condition. **G:** Schematic of the two-projection computational model. A Ca^2+^ event in Projection 1 (P1; pink) activates TRPM4, generating membrane depolarization that propagates to Projection 2 (P2; orange) to activate VGCCs. Model outputs include Δ[Ca^2+^]_i_ and TRPM4 conductance in P1, and VGCC conductance and Δ[Ca^2+^]_I_ in P2. **H:** Modeled intracellular Ca^2+^concentration in P1 over time for event amplitudes of 0.6 µM (green), 1.0 µM (pink), 2.0 µM (orange), and 10 µM (purple). **I:** Modeled TRPM4 conductance in P1 over time for the same event amplitudes. **J:** Modeled VGCC conductance in P2 over time for the same event amplitudes. **K:** Modeled intracellular Ca^2+^ concentration in P2 over time for the same event amplitudes.

**Fig. 4.**
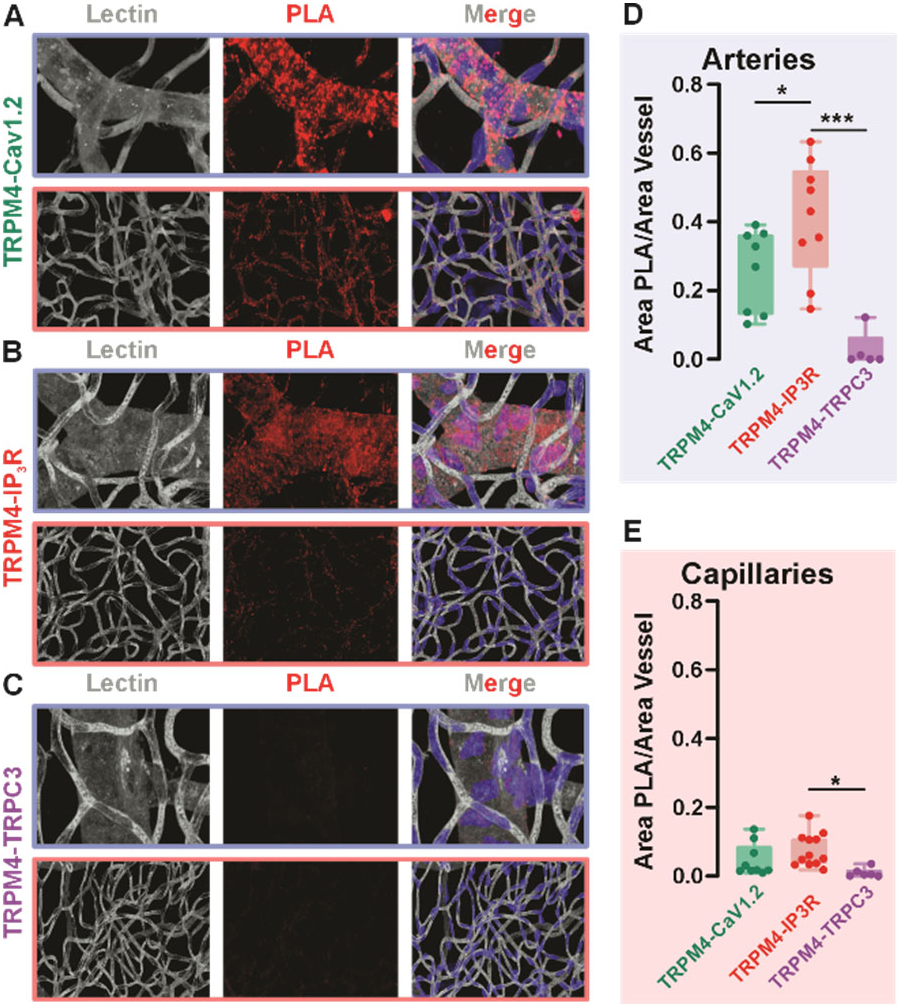
TRPM4 colocalizes with IP3 receptors at the nanoscale in retinal pericytes. **A–C:** Representative confocal images of proximity ligation assay (PLA) signal in retinal whole-mount preparations. Each row shows lectin labeling (gray, left), PLA signal (red puncta, middle), and merged images (right) for TRPM4–Cav1.2 **(A)**, TRPM4–IP_3_R **(B)**, and TRPM4–TRPC3 **(C)** protein pairs. For each pair, images from arterioles (blue border, top) and capillaries (pink border, bottom) are shown. PLA puncta indicate protein pairs within ≤40 nm. Lectin labeling provides anatomical context for vascular localization of PLA signal. **D:** Quantification of PLA signal normalized to vessel area (Area PLA/Area Vessel) for each protein pair in arterioles. TRPM4–IP_3_R proximity was significantly greater than both TRPM4–Cav1.2 and TRPM4–TRPC3, indicating preferential nanoscale colocalization of TRPM4 and IP_3_R in arteriolar mural cells. **E:** Quantification of PLA signal normalized to vessel area for each protein pair in capillaries. TRPM4–IP_3_R proximity was significantly greater than TRPM4–TRPC3, confirming preferential nanoscale association of TRPM4 and IP_3_R in capillary pericytes. Data are presented as median with interquartile range. Scale bar, 10µm. Comparisons across protein pairs were made using one-way ANOVA with Tukey’s post hoc test. *P ≤ 0.05, ***P ≤ 0.001; n = 7-9 preparations from N = 3 mice per condition.

With two kinetically and mechanistically distinct Ca^2+^ event populations, we next asked whether these differences in Ca^2+^ signal shape have functional consequences for inter-projection signaling. TRPM4 channels exhibits a characteristic bell-shaped dependence on intracellular Ca^2+^ concentration^14, 37^, making its activation highly sensitive to both the amplitude and duration of Ca^2+^ signals. We therefore reasoned that the slow, sustained Ca^2+^ events driven by IP_3_R activation and the fast, transient events driven by VGCC Ca^2+^ influx may differ in their capacity to activate TRPM4, generate membrane depolarization, and propagate electrical signals to VGCCs in a neighboring projection. To test this, we developed a computational model incorporating the Ca^2+^ event kinetics characterized above as inputs to simulate TRPM4 activation, membrane depolarization, and downstream VGCC activation in a distal pericyte projection (Fig. 3G). This model was constructed around the essential signaling components required to simulate Ca^2+^-dependent pericyte contraction. To remain within physiological limits, the maximal intracellular Ca^2+^ concentration was constrained to 1 mM^40, 57^. TRPM4 conductance was represented using a Gaussian response function, capturing its bell-shaped dependence on intracellular Ca^2+14, 37^, whereas VGCC activation was described by a Boltzmann equation to model voltage-dependent gating dynamics^35^. A Ca^2+^ buffering mechanism was incorporated to maintain intracellular concentrations within physiological ranges^57^. We limited the model to signaling elements previously characterized in pericytes and vascular smooth muscle cells^8, 45^. Although the underlying biology is substantially more complex, this simplified, equation-based framework provides a tractable foundation for investigating projection-specific signaling and ion channel contributions, with the flexibility for future refinement as additional mechanisms are defined.

To model biologically relevant Ca^2+^ events in the initiating projection (P1), we reproduced the traces presented in Fig. 3C. At 0.6 and 1 µM [Ca^2+^], signals were characterized by a gradual rise, sustained plateau, and slow decay kinetics, consistent with IP_3_R-mediated Ca^2+^ events observed in our recordings. Higher amplitude inputs (2 and 10 µM [Ca^2+^]) displayed faster rise times and more transient peaks (Fig. 3H), consistent with rapid, high-amplitude Ca^2+^ influx from VGCCs observed *ex vivo* (Fig. 3C).

We then investigated how these differences in Ca^2+^ signal shape in P1 affect TRPM4 recruitment and signal propagation in the distal projection (P2). We examined how shape of the Ca^2+^ signals influence TRPM4 activation. At 0.6 µM [Ca^2+^], TRPM4 exhibited a prolonged activation profile providing a consistent depolarizing current (Fig. 3I). Higher concentration inputs triggered rapid but short-lived TRPM4 activation, indicating that sharp Ca^2+^ spikes drive quick channel inactivation (Fig. 3I). To further characterize the concentration-dependence of TRPM4 activation across a range of physiologically relevant Ca^2+^ amplitudes, we systematically varied input concentrations. At low-to-moderate Ca^2+^ amplitudes (0.5–0.7 µM), TRPM4 conductance profiles were similar to the 0.6 µM condition, with sustained activation (Sup. Figure 5). At moderate-to-high amplitudes (0.5–1.5 µM), a progressive transition from sustained to more transient TRPM4 activation was observed (Sup. Figure 6). This transition became more pronounced at higher amplitudes (0.5–2.0 µM), where TRPM4 activation was increasingly brief (Sup. Figure 7). At very high amplitudes (0.5–3.0 µM), TRPM4 activation was further curtailed, consistent with rapid channel inactivation (Sup. Figure 8). We next modeled VGCC conductance over time in response to TRPM4-driven membrane depolarization to determine whether Ca^2+^ signals originating in P1 can initiate Ca^2+^ influx in P2. At 0.6 µM [Ca^2+^], a gradual, sustained increase in VGCC conductance reached a maximum of 60 pS, effectively promoting Ca^2+^ influx (Fig. 3J). At higher concentrations (1.0–10 µM), VGCC conductance increased only transiently, failing to sustain activation, likely due to premature channel inactivation or insufficient dwell time near the activation threshold (Fig. 3J). At extreme Ca^2+^ amplitudes (10–25 µM), TRPM4 activation and downstream VGCC conductance were markedly diminished, further supporting the conclusion that supraphysiological Ca^2+^ inputs are ineffective at driving sustained signal propagation (Sup. Figure 9).

**Figure. 5.**
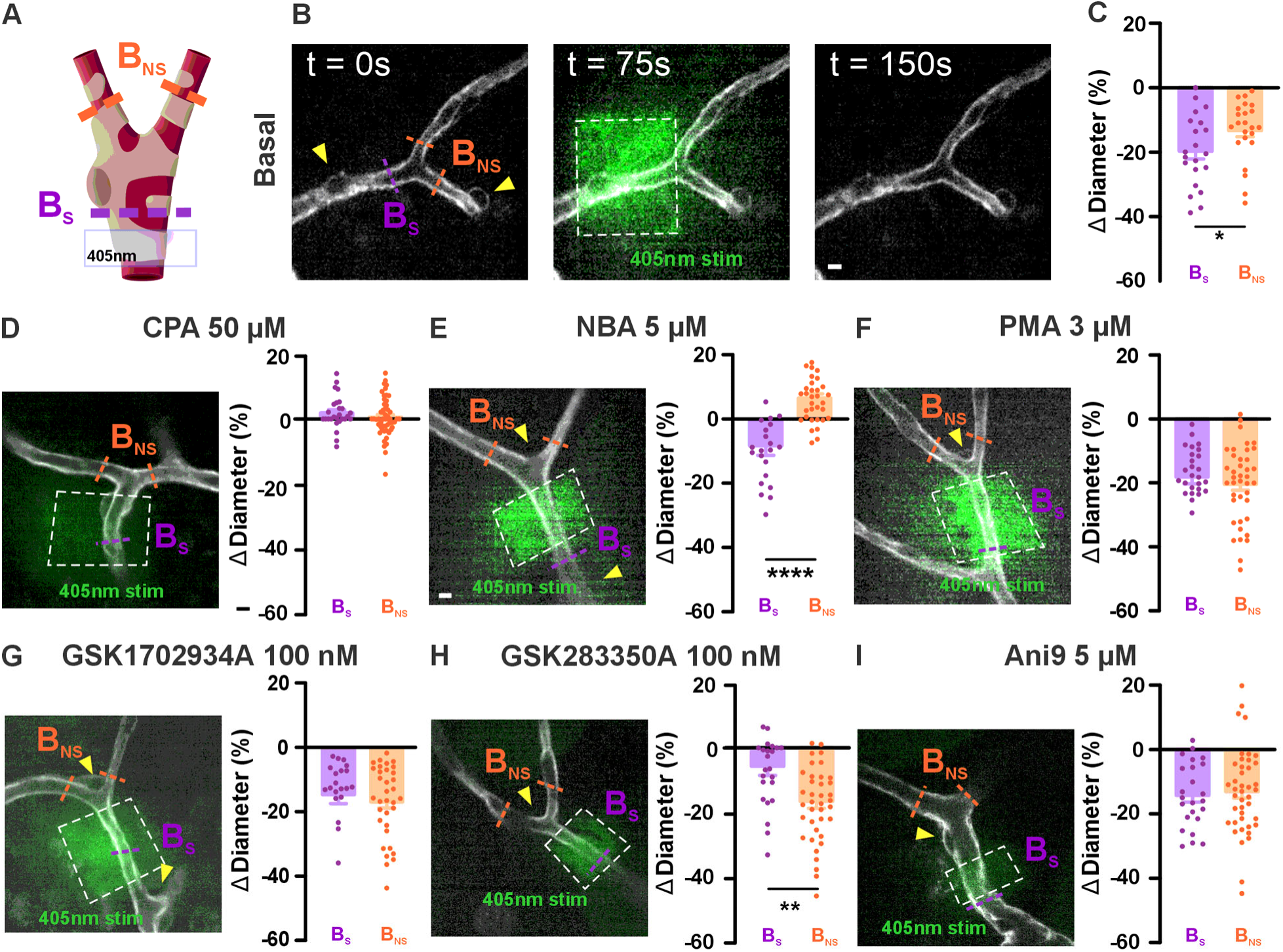
Local IP_3_ photolysis activates TRPM4-dependent projection-to-projection capillary constriction in retinal pericytes. **A:** Schematic illustrating the experimental design. Focal 405 nm illumination (gray box) was targeted to the stimulated branch (B_S_, purple) at a capillary bifurcation, while diameter changes were simultaneously monitored in the non-stimulated branch (B_NS_, orange). **B:** Representative confocal images at baseline (t = 0 s), during stimulation (t = 75 s), and post-stimulation (t = 150 s) under basal conditions. B_S_ (purple) and B_NS_ (orange) are indicated. Yellow arrowheads denote location of pericyte nucleus. Scale bar, 10 µm. **C:** Summary data of percent diameter change (ΔDiameter, %) in B_S_ (purple) and B_NS_ (orange) under basal conditions. **(D–I)** Representative confocal images during 405 nm stimulation and summary data of ΔDiameter in B_S_ and B_NS_ following bath application of CPA (50 µM; D), NBA (5 µM; E), PMA (3 µM; F), GSK1702934A (100 nM; G), GSK2833503A (100 nM; H), or Ani9 (5 µM; I). Data are presented as means ± SEM. Comparisons between B_S_ and B_NS_ within each condition were made using unpaired t-tests. *P ≤ 0.05, ****P ≤ 0.0001 versus indicated comparison; n = 7-9 capillaries per group from N = 3 mice per condition.

Together, these results suggest that sustained, low-amplitude Ca^2+^ signals are more effective at maintaining TRPM4 activation than brief, high-amplitude transients. Finally, we asked whether TRPM4-induced depolarization and VGCC activation are sufficient to drive Ca^2+^ influx into P2. Using the VGCC conductance profiles from Fig. 3J, we modeled Ca^2+^ concentration in P2 over time (Fig. 3K). The 0.6 µM input generated the largest and most stable Ca^2+^ entry into P2. In contrast, higher concentration inputs in P1, which failed to sustain TRPM4 or VGCC activity, produced diminished Ca^2+^ influx in P2. Collectively, these simulations demonstrate that Ca^2+^ event kinetics, rather than amplitude alone, determine the efficacy of inter-projection signal propagation, with sustained, low-amplitude IP_3_R-mediated Ca^2+^ events preferentially activating TRPM4, depolarizing the membrane, and driving VGCC-mediated Ca^2+^ influx in distal projections, consistent with the Ca^2+^-dependent gating properties of TRPM4 in other cell types^16, 58–60^.

### TRPM4 is in close proximity to IP_3_ receptors in pericytes

Our simulations predict that sustained, low-amplitude IP_3_R-mediated Ca^2+^ events preferentially activate TRPM4 to drive inter-projection signal propagation, implying that IP_3_Rs and TRPM4 must be in sufficient proximity to allow local Ca^2+^ transfer between the two channels. To test this, we used the proximity ligation assay (PLA) to determine whether IP_3_ receptors and TRPM4 channels are spatially organized to support localized Ca^2+^-dependent activation in pericytes. PLA detects protein pairs within 10 nm of one another, making it a powerful tool for identifying functionally relevant molecular interactions *in situ* without requiring co-immunoprecipitation or overexpression systems. We tested the proximity of TRPM4 with Cav1.2, IP_3_ receptors (IP_3_R), and TRPC3. Lectin staining was used to outline the vasculature and distinguish smooth muscle cell (SMC) from pericyte coverage. TRPM4-Cav1.2 PLA signal (Fig. 4A) was concentrated in arteriolar regions, consistent with SMC localization, with sparse signal at the capillary level where pericytes reside and minimal signal in venules where SMC coverage is absent (Fig. 4D–F). TRPM4-IP_3_R PLA signal was detected throughout the vascular tree (Fig. 4B), with a higher proportion in arterioles, likely reflecting greater SMC density relative to pericytes (Fig. 4D–F). In contrast, TRPM4-TRPC3 PLA signal was largely absent across the retinal vasculature (Fig. 4C, D–F), suggesting these channels do not interact in close proximity. Together, these findings indicate that IP_3_R is a privileged molecular partner for TRPM4 in the retinal vasculature and support a mechanism whereby IP_3_R-mediated Ca^2+^ release drives local TRPM4 activation in pericytes.

### Local IP_3_ Uncaging Activates TRPM4-Mediated Projection-to-Projection Constriction

Our computational model predicted that sustained IP_3_R-mediated Ca^2+^release activates TRPM4 within a stimulated pericyte projection, generating membrane depolarization that propagates to distal projections and activates voltage-gated Ca^2+^ channels (VGCCs). To experimentally test this mechanism, we performed focal photolysis of caged IP_3_ in flat-mounted retina preparations, enabling spatially restricted IP_3_R-mediated Ca^2+^ release within individual pericyte projections (Fig. 5A). Localized 405 nm stimulation produced robust constriction at the stimulated branch (B_S_, −19.64%) and a smaller but significant constriction at the non-stimulated branch (B_NS_, −13.21%; Fig. 5B and C), indicating that local IP_3_R activation generates a signal capable of propagating between projections to drive distal constriction. To determine whether intracellular Ca^2+^ store release is required for this response, retinas were treated with cyclopiazonic acid (CPA; 50 µM) to inhibit SERCA-dependent Ca^2+^ store refilling. Under these conditions, IP_3_ uncaging failed to evoke constriction in either the stimulated or non-stimulated projection (Fig. 5D), demonstrating that IP_3_-mediated constriction and the projection-to-projection signaling requires intracellular Ca^2+^ store mobilization. We next tested whether TRPM4 mediates the propagation of this signal by inhibiting the channel with NBA (5 µM). Under these conditions, IP_3_ uncaging produced only modest constriction at the stimulated projection (−8.00%), while the non-stimulated projection failed to constrict and instead exhibited a small dilation (+3.05%; Fig. 5E). These findings suggest that TRPM4 activity is required for propagation of the depolarizing signal to distal projections. Because TRPM4 activity is enhanced by PKC-dependent membrane trafficking^53^, we next activated PKC with PMA (3 µM) to increase surface expression of TRPM4. PMA treatment enhanced constriction at both the stimulated (−18.11%) and non-stimulated (−20.34%) projections (Fig. 5F), consistent with amplified depolarization-dependent Ca^2+^ influx throughout the projection network. To further examine the contribution of depolarizing cation conductances to projection-to-projection signaling, we activated TRPC3 with GSK1702934A (100 nM). TRPC3 activation enhanced constriction responses following IP_3_ uncaging, particularly within distal projections (16.75%, Fig. 5G), supporting the idea that increased depolarizing cation influx can facilitate projection-to-projection signaling between pericyte projections. Conversely, inhibition of TRPC3 with GSK2833503A (100 nM) preferentially attenuated constriction within the stimulated projection while largely preserving distal constriction (Fig. 5H), suggesting that TRPC3 primarily contributes to local depolarization and constriction rather than long-range signal propagation. In contrast, inhibition of Ano1 with Ani9 (5 µM) produced comparatively modest effects on propagated constriction (Fig. 5I), indicating that Ano1 plays a limited role in projection-to-projection signaling under these conditions. Together, these findings support a model in which localized IP_3_R-mediated Ca^2+^ release activates TRPM4 within stimulated pericyte projections, generating a propagating depolarization that recruits voltage-dependent Ca^2+^ influx and coordinated constriction in distal projections. While TRPC3-dependent cation influx appears to augment local constriction responses, TRPM4-mediated depolarization likely serves as the primary driver of long-range signaling across pericyte projection networks.

## Discussion

The present study identifies TRPM4 as a critical mediator of electrical coupling and signal propagation between ensheathing pericyte projections. Using a combination of optogenetic stimulation, computational modeling, proximity ligation, and *ex vivo* pharmacology, we demonstrate that pericyte projections operate as semi-autonomous contractile units capable of both spatially confined and propagated signaling. Our findings reveal that GPCR-mediated second messenger signaling remains confined to the stimulated projection, whereas membrane depolarization propagates to neighboring projections via TRPM4, enabling coordinated constriction across multiple capillary branches. Together, these results establish a mechanistic framework for projection-specific control of microvascular perfusion.

### Pressure-Induced Tone and Physiological Relevance of Pericyte-Mediated Microvascular Control

A defining feature of the microcirculation is its capacity to autoregulate blood flow in response to changes in intraluminal pressure, classically attributed to the myogenic response of vascular smooth muscle cells in arterioles^13, 50, 54, 61^. Most of our mechanistic insight into pressure-induced vasoconstriction comes from studies of linear vessels such as arterioles, where smooth muscle cells wrap circumferentially around the vessel wall, sense wall tension as a single functional unit, and generate coordinated contractile force to resist pressure-induced distension. However, the capillary network is fundamentally different in architecture: rather than linear tubes, capillaries form complex bifurcating networks in which individual branches vary in diameter, pressure, and flow. At these bifurcation points, ensheathing pericytes extend multiple projections that wrap around capillary branches of different diameters, each of which constitutes an independent mechanosensory and contractile domain^5^. Consistent with Laplace’s law, pericyte projections distributed across branches of different diameters are exposed to varying degrees of circumferential stretch and wall tension. Unlike smooth muscle cells, which integrate tension across the entire cell, pericytes must therefore integrate mechanosensory information across spatially distinct projections operating under different tensional loads. Our findings suggest that TRPM4 serves as a critical node for this integration. Pressure-induced GqPCR activation and downstream IP_3_/DAG generation trigger localized Ca^2+^ release in individual projections, activating TRPM4 and generating membrane depolarization that can propagate electrically to neighboring projections to coordinate contractile responses at the capillary bifurcation. In this way, the pericyte-TRPM4 signaling axis represents a fundamentally more sophisticated pressure-sensing mechanism than the smooth muscle myogenic response. Whereas a smooth muscle cell integrates a single mechanical input from one vessel and generates a uniform contractile response, an ensheathing pericyte simultaneously samples hemodynamic conditions across multiple capillary branches, integrates these heterogeneous tensional signals through TRPM4-dependent electrical coupling, and generates a spatially coordinated contractile output across the bifurcating network. This capacity for multi-branch mechanical integration would endow the capillary network with a level of local flow regulation far beyond what a simple myogenic response could achieve, enabling dynamic, branch-specific control of blood flow distribution at the finest level of the vasculature.

### Compartmentalization of Ca^2+^ and Membrane Potential in Pericyte Projections: A Stimulus-Dependent Molecular Switch

A central conceptual contribution of this study is the demonstration that ensheathing pericyte projections operate as semi-autonomous signaling domains capable of independently compartmentalizing both Ca^2+^ signals and membrane potential, a principle that distinguishes pericytes fundamentally from vascular smooth muscle cells and has important implications for how capillary blood flow is regulated. SMCs are elongated, circumferentially oriented cells that wrap uniformly around arteriolar walls, and their contractile responses are coordinated through gap junction-mediated electrical coupling and propagating Ca^2+^ waves that synchronize tone across large vascular segments^10, 62^. This uniform electrical organization is well-suited to the function of arterioles as pressure-regulating conduits, but it precludes the kind of branch-specific, spatially graded responses that capillary flow regulation requires. In contrast, pericyte projections are morphologically and functionally heterogeneous, extending along and around individual capillary branches, and our data demonstrate that their contractile responses can be branch-specific, supporting the emerging view that pericytes are active participants in microvascular flow control rather than passive structural elements^63, 64^.

This organizational principle has important parallels in neurons and astrocytes, where the subcellular segregation of both biochemical and electrical signals underlies the remarkable computational capacity of these cells. In neurons, individual dendritic branches can generate branch-specific Ca^2+^ transients and sustain local membrane potential gradients that differ substantially from those at the soma, arising from the localized distribution of voltage-gated and ligand-gated ion channels combined with the spatial restriction of second messenger cascades^33^. Similarly, astrocytic processes maintain spatially distinct membrane potential microdomains across their morphologically complex arbors, enabling local responses to synaptic activity without necessarily engaging the entire cell^65^. Our findings suggest that pericyte projections exploit an analogous architecture: Ca^2+^ signals can remain confined to individual projections or propagate to neighboring ones depending on the nature of the initiating stimulus, and membrane depolarization generated in one projection can propagate electrically to activate VGCCs in a distal projection via TRPM4. We therefore propose that the subcellular compartmentalization of Ca^2+^ and electrical signaling in pericyte projections represents a specialized adaptation for fine-grained, branch-specific regulation of capillary perfusion, a function that is neither required nor present in the more uniform contractile machinery of vascular smooth muscle. Critically, this implies that individual pericyte projections may maintain distinct membrane potentials simultaneously, a form of electrical compartmentalization not previously described in vascular mural cells and representing a fundamental departure from the uniform electrical coupling of vascular smooth muscle. Resolving this principle will require electrophysiological or optical approaches capable of measuring subcellular voltage dynamics at the level of individual projections, an important direction for future investigation.

### Context-Dependent Signaling Determines the Spatial Boundary of Pericyte Contraction

The apparent contradiction between our opto-α1AR and IP_3_ uncaging experiments is central to understanding pericyte projection compartmentalization. Opto-α1AR stimulation simultaneously generates IP_3_ and DAG yet produces branch-specific constriction without engaging neighboring projections, whereas selective IP_3_ photolysis, without concurrent DAG generation, produced constriction at both stimulated and distal projections in a TRPM4-dependent manner. The resolution of this apparent contradiction lies not in direct antagonism between TRPC3 and TRPM4, but in how concurrent DAG signaling alters Ca^2+^ signal kinetics. During opto-α1AR stimulation, DAG-mediated TRPC3 activation drives rapid Ca^2+^ influx that augments local contraction but generates a Ca^2+^ signal too brief and high-amplitude to sustain the prolonged, low-amplitude dynamics required for optimal TRPM4 activation, consistent with the bell-shaped Ca^2+^ dependence of TRPM4 and our computational modeling. In support of this, pharmacological TRPC3 activation during IP_3_ uncaging enhanced distal projection constriction, while TRPC3 inhibition selectively attenuated local constriction while preserving distal responses, confirming that TRPC3 functions primarily as a local amplifier of contractile force rather than a suppressor of inter-projection signaling, consistent with the recently identified role of TRPC3 as the primary depolarizing mechanism driving pressure-induced constriction in transitional pericytes of the brain microcirculation^6^. Together, these findings indicate that the nature of the vasoactive stimulus, through its differential engagement of IP_3_ and DAG pathways, acts as a molecular switch determining the spatial reach of the pericyte contractile response, mirroring the conditional signal integration described in neurons.

### Ion Channels Operating in Series, Parallel, and Opposition Across Pericyte Projections

A further consequence of projection-level compartmentalization is that ion channels within and between projections can interact in series, in parallel, or in opposition. Although such channel interactions have been extensively studied in vascular smooth muscle cells, they are often examined at the level of individual channels or signaling pathways, with less consideration given to how subcellular compartmentalization shapes their integrated behavior. Within a single projection, IP_3_R-mediated Ca^2+^ release acts in series with TRPM4 activation and sequential VGCC opening, forming a signal amplification cascade that converts a local biochemical input into a graded contractile output. Across projections, TRPM4-mediated electrical coupling allows VGCCs in distal projections to operate in parallel with those in the stimulated projection, extending a branch-specific response into a coordinated, network-level contractile event. The capacity to switch between these two modes, determined by the molecular switch described above, provides the pericyte with a versatile range of hemodynamic outputs from a common signaling architecture. Perhaps most intriguingly, TRPC3 and TRPM4 operate in functional opposition: DAG-mediated TRPC3 activation generates rapid local depolarization and contraction while simultaneously limiting the sustained Ca^2+^ signal required for TRPM4-dependent inter-projection propagation, effectively gating the spatial boundary of the contractile response. This antagonistic interaction between channels sharing a common upstream activator mirrors the competing conductances that shape synaptic signal integration in neurons, but represents a previously unrecognized principle in pericyte physiology. Together, series amplification within projections, parallel activation across projections, and opposing channel interactions at the propagation boundary, these organizational principles establish pericyte projections as computationally sophisticated signaling units capable of precise, context-dependent regulation of capillary perfusion.

### The Pathophysiological Implications of this Level of Cellular Integration are Significant

Pericyte loss or dysfunction has been implicated in a broad spectrum of neurological and vascular diseases, including epilepsy, multiple sclerosis, hypertension, Alzheimer’s disease, ischemia-reperfusion injury, and diabetic retinopathy^63, 64^. A critical but underappreciated consequence of pericyte loss may not be the complete failure of blood flow regulation, but rather the degradation of its precision and efficiency. In the absence of active pericyte-mediated tone, blood flow distribution across the capillary network would revert to passive governance determined solely by the fixed geometric properties of the network, like vessel radius, length, and branching architecture, rather than by dynamic, activity-dependent adjustments. In support of this, *in vivo* suppression of TRPM4 in cerebral arteries impaired autoregulatory vasoconstriction and significantly elevated cerebral blood flow across a range of arterial pressures, demonstrating that loss of TRPM4-dependent tone compromises the precision of blood flow regulation rather than eliminating it entirely^16, 59, 66, 67^. In the brain and retina, where neural activity is spatially and temporally heterogeneous, this loss of regulatory precision would be particularly consequential, reducing the capillary network from an actively regulated, demand-responsive system to a passively perfused one, insufficient to meet the dynamic metabolic demands of neural activity. Notably, while TRPM4 knockout mice do not exhibit an overt baseline neurological phenotype^68^, likely reflecting compensatory adaptations, they show significantly improved cognitive outcomes and reduced neuronal injury following status epilepticus^69^, suggesting that TRPM4 activity becomes critically detrimental when blood flow regulation is challenged under pathological conditions. These findings position TRPM4 not merely as a regulator of capillary tone, but as a critical determinant of neurovascular efficiency, identifying it as a compelling therapeutic target for preserving microvascular function across a spectrum of retinal and brain diseases.

## Conclusion

The present study establishes TRPM4 as a critical mediator of electrical coupling between ensheathing pericyte projections, enabling dynamic, branch-specific regulation of capillary perfusion. The spatial reach of pericyte contractile responses is determined by the nature of the initiating signal: sustained IP_3_R-mediated Ca^2+^ release propagates depolarization and coordinated constriction across distal projections via TRPM4, while concurrent DAG signaling confines responses to the stimulated branch. This stimulus-dependent gating, supported by the nanoscale organization of IP_3_R and TRPM4 at the pericyte membrane, positions ensheathing pericytes as computationally sophisticated signaling units capable of context-dependent microvascular flow control. How this mechanism is modulated by vasoactive signals, neural activity, and pathological conditions such as diabetic retinopathy and cerebrovascular disease remains an important direction for future investigation.

## Acknowledgments

This study was supported by grants from the NIH (NIA R01AG081935, NHLBI R01HL177084, K01HL138215, and NIGMS P20GM130459 to A.L.G.). The Transgenic Genotyping and Phenotyping Core and the High Spatial and Temporal Resolution Imaging Core at the Centers of Biomedical Research Excellence (COBRE) Center for Molecular and Cellular Signaling in the Cardiovascular System, University of Nevada, Reno are maintained by grants from NIH/NIGMS (P20GM130459 Sub#5451 and P20GM130459 Sub#5452). We thank C. Franco, E. Franco, P. Voss, and M. Dagda for technical assistance and animal care assistance. We also thank M.T. Nelson for his contributions to the original Ca²⁺ imaging dataset re-analyzed in this study.

## Author contributions

V.M., A.A, and A.L.G. designed research; A.A, A.E, and A.L.G. performed experiments; V.M performed computational modeling; V.M., A.A, and A.L.G. analyzed data; and V.M., A.A, and A.L.G. wrote the paper.

## References

1. Mughal A, Nelson MT, Hill-Eubanks D. The post-arteriole transitional zone: a specialized capillary region that regulates blood flow within the CNS microvasculature. J Physiol. 2023;601(5):889–901. Epub 20230221. doi: 10.1113/JP282246. PubMed PMID: 36751860; PMCID: PMC9992301.

2. Hartmann DA, Underly RG, Grant RI, Watson AN, Lindner V, Shih AY. Pericyte structure and distribution in the cerebral cortex revealed by high-resolution imaging of transgenic mice. Neurophotonics. 2015;2(4):041402. Epub 20150527. doi: 10.1117/1.NPh.2.4.041402. PubMed PMID: 26158016; PMCID: PMC4478963.

3. Grant RI, Hartmann DA, Underly RG, Berthiaume AA, Bhat NR, Shih AY. Organizational hierarchy and structural diversity of microvascular pericytes in adult mouse cortex. J Cereb Blood Flow Metab. 2019;39(3):411–25. Epub 2017/09/22. doi: 10.1177/0271678X17732229. PubMed PMID: 28933255; PMCID: PMC6399730.

4. Fernandez-Klett F, Offenhauser N, Dirnagl U, Priller J, Lindauer U. Pericytes in capillaries are contractile in vivo, but arterioles mediate functional hyperemia in the mouse brain. Proc Natl Acad Sci U S A. 2010;107(51):22290–5. Epub 2010/12/08. doi: 10.1073/pnas.1011321108. PubMed PMID: 21135230; PMCID: PMC3009761.

5. Gonzales AL, Klug NR, Moshkforoush A, Lee JC, Lee FK, Shui B, Tsoukias NM, Kotlikoff MI, Hill-Eubanks D, Nelson MT. Contractile pericytes determine the direction of blood flow at capillary junctions. Proc Natl Acad Sci U S A. 2020;117(43):27022–33. Epub 20201013. doi: 10.1073/pnas.1922755117. PubMed PMID: 33051294; PMCID: PMC7604512.

6. Ferris HR, Jeffrey DA, Guerrero MB, Birnbaumer L, Zheng F, Dabertrand F. Increased luminal pressure in brain capillaries drives TRPC3-dependent depolarization and constriction of transitional pericytes. Sci Signal. 2025;18(884):eads1903. Epub 20250429. doi: 10.1126/scisignal.ads1903. PubMed PMID: 40299956.

7. Klug NR, Sancho M, Gonzales AL, Heppner TJ, O’Brien RIC, Hill-Eubanks D, Nelson MT. Intraluminal pressure elevates intracellular calcium and contracts CNS pericytes: Role of voltage-dependent calcium channels. Proc Natl Acad Sci U S A. 2023;120(9):e2216421120. Epub 20230221. doi: 10.1073/pnas.2216421120. PubMed PMID: 36802432; PMCID: PMC9992766.

8. Hill-Eubanks DC, Werner ME, Heppner TJ, Nelson MT. Calcium signaling in smooth muscle. Cold Spring Harb Perspect Biol. 2011;3(9):a004549. Epub 20110901. doi: 10.1101/cshperspect.a004549. PubMed PMID: 21709182; PMCID: PMC3181028.

9. Nelson MT, Patlak JB, Worley JF, Standen NB. Calcium channels, potassium channels, and voltage dependence of arterial smooth muscle tone. Am J Physiol. 1990;259(1 Pt 1):C3-18. doi: 10.1152/ajpcell.1990.259.1.C3. PubMed PMID: 2164782.

10. Somlyo AP, Somlyo AV. Signal transduction and regulation in smooth muscle. Nature. 1994;372(6503):231–6. doi: 10.1038/372231a0. PubMed PMID: 7969467.

11. Burdyga T, Borysova L. Calcium signalling in pericytes. J Vasc Res. 2014;51(3):190–9. Epub 20140604. doi: 10.1159/000362687. PubMed PMID: 24903335.

12. Hofmann T, Obukhov AG, Schaefer M, Harteneck C, Gudermann T, Schultz G. Direct activation of human TRPC6 and TRPC3 channels by diacylglycerol. Nature. 1999;397(6716):259–63. doi: 10.1038/16711. PubMed PMID: 9930701.

13. Earley S, Brayden JE. Transient receptor potential channels in the vasculature. Physiol Rev. 2015;95(2):645–90. doi: 10.1152/physrev.00026.2014. PubMed PMID: 25834234; PMCID: PMC4551213.

14. Launay P, Fleig A, Perraud AL, Scharenberg AM, Penner R, Kinet JP. TRPM4 is a Ca2+-activated nonselective cation channel mediating cell membrane depolarization. Cell. 2002;109(3):397–407. doi: 10.1016/s0092-8674(02)00719-5. PubMed PMID: 12015988.

15. Abriel H, Syam N, Sottas V, Amarouch MY, Rougier JS. TRPM4 channels in the cardiovascular system: physiology, pathophysiology, and pharmacology. Biochem Pharmacol. 2012;84(7):873–81. Epub 20120627. doi: 10.1016/j.bcp.2012.06.021. PubMed PMID: 22750058.

16. Earley S, Waldron BJ, Brayden JE. Critical role for transient receptor potential channel TRPM4 in myogenic constriction of cerebral arteries. Circ Res. 2004;95(9):922–9. Epub 20041007. doi: 10.1161/01.RES.0000147311.54833.03. PubMed PMID: 15472118.

17. Shui B, Lee JC, Reining S, Lee FK, Kotlikoff MI. Optogenetic sensors and effectors: CHROMus-the Cornell Heart Lung Blood Institute Resource for Optogenetic Mouse Signaling. Front Physiol. 2014;5:428. Epub 2014/11/22. doi: 10.3389/fphys.2014.00428. PubMed PMID: 25414670; PMCID: PMC4222331.

18. Harris CR, Millman KJ, van der Walt SJ, Gommers R, Virtanen P, Cournapeau D, Wieser E, Taylor J, Berg S, Smith NJ, Kern R, Picus M, Hoyer S, van Kerkwijk MH, Brett M, Haldane A, Del Rio JF, Wiebe M, Peterson P, Gerard-Marchant P, Sheppard K, Reddy T, Weckesser W, Abbasi H, Gohlke C, Oliphant TE. Array programming with NumPy. Nature. 2020;585(7825):357–62. Epub 20200916. doi: 10.1038/s41586-020-2649-2. PubMed PMID: 32939066; PMCID: PMC7759461.

19. Hunter JD. Matplotlib: A 2D Graphics Environment. Computing in Science & Engineering. 2007;vol. 9, no. 3,: pp. 90–5. doi: 10.1109/MCSE.2007.55.

20. Destexhe A, Huguenard JR. Nonlinear thermodynamic models of voltage-dependent currents. J Comput Neurosci. 2000;9(3):259–70. doi: 10.1023/a:1026535704537. PubMed PMID: 11139042.

21. Vennekens R, Nilius B. Insights into TRPM4 function, regulation and physiological role. Handb Exp Pharmacol. 2007(179):269–85. doi: 10.1007/978-3-540-34891-7_16. PubMed PMID: 17217063.

22. Dietrich A, Mederos YSM, Gollasch M, Gross V, Storch U, Dubrovska G, Obst M, Yildirim E, Salanova B, Kalwa H, Essin K, Pinkenburg O, Luft FC, Gudermann T, Birnbaumer L. Increased vascular smooth muscle contractility in TRPC6-/- mice. Mol Cell Biol. 2005;25(16):6980–9. doi: 10.1128/MCB.25.16.6980-6989.2005. PubMed PMID: 16055711; PMCID: PMC1190236.

23. Caputo A, Caci E, Ferrera L, Pedemonte N, Barsanti C, Sondo E, Pfeffer U, Ravazzolo R, Zegarra-Moran O, Galietta LJ. TMEM16A, a membrane protein associated with calcium-dependent chloride channel activity. Science. 2008;322(5901):590–4. Epub 20080904. doi: 10.1126/science.1163518. PubMed PMID: 18772398.

24. Hartzell HC, Whitlock JM. TMEM16 chloride channels are two-faced. J Gen Physiol. 2016;148(5):367–73. doi: 10.1085/jgp.201611686. PubMed PMID: 27799317; PMCID: PMC5089938.

25. Korte N, Ilkan Z, Pearson CL, Pfeiffer T, Singhal P, Rock JR, Sethi H, Gill D, Attwell D, Tammaro P. The Ca2+-gated channel TMEM16A amplifies capillary pericyte contraction and reduces cerebral blood flow after ischemia. J Clin Invest. 2022;132(9). doi: 10.1172/JCI154118. PubMed PMID: 35316222; PMCID: PMC9057602.

26. Davey MJ. The pharmacology of prazosin, an alpha 1-adrenoceptor antagonist and the basis for its use in the treatment of essential hypertension. Clin Exp Hypertens A. 1982;4(1-2):47–59. doi: 10.3109/10641968209061575. PubMed PMID: 6122521.

27. Seidler NW, Jona I, Vegh M, Martonosi A. Cyclopiazonic acid is a specific inhibitor of the Ca2+-ATPase of sarcoplasmic reticulum. J Biol Chem. 1989;264(30):17816–23. PubMed PMID: 2530215.

28. Kazda S, Towart R. Nimodipine: a new calcium antagonistic drug with a preferential cerebrovascular action. Acta Neurochir (Wien). 1982;63(1-4):259–65. doi: 10.1007/BF01728880. PubMed PMID: 7102417.

29. Kiyonaka S, Kato K, Nishida M, Mio K, Numaga T, Sawaguchi Y, Yoshida T, Wakamori M, Mori E, Numata T, Ishii M, Takemoto H, Ojida A, Watanabe K, Uemura A, Kurose H, Morii T, Kobayashi T, Sato Y, Sato C, Hamachi I, Mori Y. Selective and direct inhibition of TRPC3 channels underlies biological activities of a pyrazole compound. Proc Natl Acad Sci U S A. 2009;106(13):5400–5. Epub 20090316. doi: 10.1073/pnas.0808793106. PubMed PMID: 19289841; PMCID: PMC2664023.

30. Kleinlogel S, Feldbauer K, Dempski RE, Fotis H, Wood PG, Bamann C, Bamberg E. Ultra light-sensitive and fast neuronal activation with the Ca(2)+-permeable channelrhodopsin CatCh. Nat Neurosci. 2011;14(4):513–8. Epub 20110313. doi: 10.1038/nn.2776. PubMed PMID: 21399632.

31. Evans WH, Boitano S. Connexin mimetic peptides: specific inhibitors of gap-junctional intercellular communication. Biochem Soc Trans. 2001;29(Pt 4):606–12. doi: 10.1042/bst0290606. PubMed PMID: 11498037.

32. Marom S. Slow changes in the availability of voltage-gated ion channels: effects on the dynamics of excitable membranes. J Membr Biol. 1998;161(2):105–13. doi: 10.1007/s002329900318. PubMed PMID: 9435267.

33. London M, Hausser M. Dendritic computation. Annu Rev Neurosci. 2005;28:503–32. doi: 10.1146/annurev.neuro.28.061604.135703. PubMed PMID: 16033324.

34. Unwin N. The structure of ion channels in membranes of excitable cells. Neuron. 1989;3(6):665–76. doi: 10.1016/0896-6273(89)90235-3. PubMed PMID: 2484344.

35. Catterall WA. Voltage-gated calcium channels. Cold Spring Harb Perspect Biol. 2011;3(8):a003947. Epub 20110801. doi: 10.1101/cshperspect.a003947. PubMed PMID: 21746798; PMCID: PMC3140680.

36. Clapham DE. Calcium Signaling. Cell. 2007;131(6):1047–58. doi: 10.1016/j.cell.2007.11.028.

37. Nilius B, Droogmans G. Ion channels and their functional role in vascular endothelium. Physiol Rev. 2001;81(4):1415–59. doi: 10.1152/physrev.2001.81.4.1415. PubMed PMID: 11581493.

38. Schuster S, Marhl M, Hofer T. Modelling of simple and complex calcium oscillations. From single-cell responses to intercellular signalling. Eur J Biochem. 2002;269(5):1333–55. doi: 10.1046/j.0014-2956.2001.02720.x. PubMed PMID: 11874447.

39. Secomb TW, Pries AR. The microcirculation: physiology at the mesoscale. J Physiol. 2011;589(Pt 5):1047–52. Epub 20110117. doi: 10.1113/jphysiol.2010.201541. PubMed PMID: 21242255; PMCID: PMC3060585.

40. Berridge MJ. Inositol trisphosphate and calcium signalling. Nature. 1993;361(6410):315-25. doi: 10.1038/361315a0. PubMed PMID: 8381210.

41. Berridge MJ, Bootman MD, Roderick HL. Calcium signalling: dynamics, homeostasis and remodelling. Nat Rev Mol Cell Biol. 2003;4(7):517–29. doi: 10.1038/nrm1155. PubMed PMID: 12838335.

42. Koch C. Biophysics of Computation: Information Processing in Single Neurons: Oxford University Press; 1998 12 Nov 2020.

43. Pries AR, Secomb TW. Structural adaptation of microvascular networks and development of hypertension. Microcirculation. 2002;9(4):305–14. doi: 10.1038/sj.mn.7800144. PubMed PMID: 12152106.

44. Uchida K. TRPM3, TRPM4, and TRPM5 as thermo-sensitive channels. J Physiol Sci. 2024;74(1):43. Epub 20240918. doi: 10.1186/s12576-024-00937-0. PubMed PMID: 39294615; PMCID: PMC11409758.

45. Hariharan A, Weir N, Robertson C, He L, Betsholtz C, Longden TA. The Ion Channel and GPCR Toolkit of Brain Capillary Pericytes. Front Cell Neurosci. 2020;14:601324. Epub 20201218. doi: 10.3389/fncel.2020.601324. PubMed PMID: 33390906; PMCID: PMC7775489.

46. Secomb TW, Hsu R, Park EY, Dewhirst MW. Green’s function methods for analysis of oxygen delivery to tissue by microvascular networks. Ann Biomed Eng. 2004;32(11):1519–29. doi: 10.1114/b:abme.0000049036.08817.44. PubMed PMID: 15636112.

47. Hodgkin AL, Katz B. The effect of sodium ions on the electrical activity of giant axon of the squid. J Physiol. 1949;108(1):37–77. doi: 10.1113/jphysiol.1949.sp004310. PubMed PMID: 18128147; PMCID: PMC1392331.

48. Armstrong CM, Hille B. Voltage-gated ion channels and electrical excitability. Neuron. 1998;20(3):371–80. doi: 10.1016/s0896-6273(00)80981-2. PubMed PMID: 9539115.

49. Vanlandewijck M, He L, Mae MA, Andrae J, Ando K, Del Gaudio F, Nahar K, Lebouvier T, Lavina B, Gouveia L, Sun Y, Raschperger E, Rasanen M, Zarb Y, Mochizuki N, Keller A, Lendahl U, Betsholtz C. A molecular atlas of cell types and zonation in the brain vasculature. Nature. 2018;554(7693):475–80. Epub 2018/02/15. doi: 10.1038/nature25739. PubMed PMID: 29443965.

50. Mederos YSM, Storch U, Gudermann T. Mechanosensitive G(q/11) Protein-Coupled Receptors Mediate Myogenic Vasoconstriction. Microcirculation. 2016;23(8):621–5. doi: 10.1111/micc.12293. PubMed PMID: 27344060.

51. Ozhathil LC, Delalande C, Bianchi B, Nemeth G, Kappel S, Thomet U, Ross-Kaschitza D, Simonin C, Rubin M, Gertsch J, Lochner M, Peinelt C, Reymond JL, Abriel H. Identification of potent and selective small molecule inhibitors of the cation channel TRPM4. Br J Pharmacol. 2018;175(12):2504–19. Epub 20180429. doi: 10.1111/bph.14220. PubMed PMID: 29579323; PMCID: PMC6002741.

52. Grand T, Demion M, Norez C, Mettey Y, Launay P, Becq F, Bois P, Guinamard R. 9-phenanthrol inhibits human TRPM4 but not TRPM5 cationic channels. Br J Pharmacol. 2008;153(8):1697–705. Epub 20080225. doi: 10.1038/bjp.2008.38. PubMed PMID: 18297105; PMCID: PMC2438271.

53. Fish RD, Sperti G, Colucci WS, Clapham DE. Phorbol ester increases the dihydropyridine-sensitive calcium conductance in a vascular smooth muscle cell line. Circ Res. 1988;62(5):1049–54. doi: 10.1161/01.res.62.5.1049. PubMed PMID: 2452033.

54. Knot HJ, Nelson MT. Regulation of arterial diameter and wall [Ca2+] in cerebral arteries of rat by membrane potential and intravascular pressure. J Physiol. 1998;508 ( Pt 1)(Pt 1):199–209. doi: 10.1111/j.1469-7793.1998.199br.x. PubMed PMID: 9490839; PMCID: PMC2230857.

55. Nilius B, Mahieu F, Prenen J, Janssens A, Owsianik G, Vennekens R, Voets T. The Ca2+-activated cation channel TRPM4 is regulated by phosphatidylinositol 4,5-biphosphate. EMBO J. 2006;25(3):467–78. Epub 20060119. doi: 10.1038/sj.emboj.7600963. PubMed PMID: 16424899; PMCID: PMC1383543.

56. Taylor CW, Tovey SC. IP(3) receptors: toward understanding their activation. Cold Spring Harb Perspect Biol. 2010;2(12):a004010. Epub 20101027. doi: 10.1101/cshperspect.a004010. PubMed PMID: 20980441; PMCID: PMC2982166.

57. Neher E, Augustine GJ. Calcium gradients and buffers in bovine chromaffin cells. J Physiol. 1992;450:273–301. doi: 10.1113/jphysiol.1992.sp019127. PubMed PMID: 1331424; PMCID: PMC1176122.

58. Gonzales AL, Earley S. Endogenous cytosolic Ca(2+) buffering is necessary for TRPM4 activity in cerebral artery smooth muscle cells. Cell Calcium. 2012;51(1):82–93. Epub 20111207. doi: 10.1016/j.ceca.2011.11.004. PubMed PMID: 22153976; PMCID: PMC3265659.

59. Gonzales AL, Earley S. Regulation of cerebral artery smooth muscle membrane potential by Ca(2)(+)-activated cation channels. Microcirculation. 2013;20(4):337–47. doi: 10.1111/micc.12023. PubMed PMID: 23116477; PMCID: PMC3573261.

60. Diszhazi G, Magyar ZE, Lisztes E, Toth-Molnar E, Nanasi PP, Vennekens R, Toth BI, Almassy J. TRPM4 links calcium signaling to membrane potential in pancreatic acinar cells. J Biol Chem. 2021;297(3):101015. Epub 20210727. doi: 10.1016/j.jbc.2021.101015. PubMed PMID: 34329682; PMCID: PMC8371206.

61. Davis MJ, Hill MA. Signaling mechanisms underlying the vascular myogenic response. Physiol Rev. 1999;79(2):387–423. doi: 10.1152/physrev.1999.79.2.387. PubMed PMID: 10221985.

62. Welsh DG, Morielli AD, Nelson MT, Brayden JE. Transient receptor potential channels regulate myogenic tone of resistance arteries. Circ Res. 2002;90(3):248–50. doi: 10.1161/hh0302.105662. PubMed PMID: 11861411.

63. Longden TA, Zhao G, Hariharan A, Lederer WJ. Pericytes and the Control of Blood Flow in Brain and Heart. Annu Rev Physiol. 2023;85:137–64. doi: 10.1146/annurev-physiol-031522-034807. PubMed PMID: 36763972; PMCID: PMC10280497.

64. Hartmann DA, Berthiaume AA, Grant RI, Harrill SA, Koski T, Tieu T, McDowell KP, Faino AV, Kelly AL, Shih AY. Brain capillary pericytes exert a substantial but slow influence on blood flow. Nat Neurosci. 2021;24(5):633–45. Epub 20210218. doi: 10.1038/s41593-020-00793-2. PubMed PMID: 33603231; PMCID: PMC8102366.

65. Shigetomi E, Bushong EA, Haustein MD, Tong X, Jackson-Weaver O, Kracun S, Xu J, Sofroniew MV, Ellisman MH, Khakh BS. Imaging calcium microdomains within entire astrocyte territories and endfeet with GCaMPs expressed using adeno-associated viruses. J Gen Physiol. 2013;141(5):633–47. Epub 20130415. doi: 10.1085/jgp.201210949. PubMed PMID: 23589582; PMCID: PMC3639581.

66. Reading SA, Brayden JE. Central role of TRPM4 channels in cerebral blood flow regulation. Stroke. 2007;38(8):2322–8. Epub 20070621. doi: 10.1161/STROKEAHA.107.483404. PubMed PMID: 17585083.

67. Gonzales AL, Garcia ZI, Amberg GC, Earley S. Pharmacological inhibition of TRPM4 hyperpolarizes vascular smooth muscle. Am J Physiol Cell Physiol. 2010;299(5):C1195–202. Epub 2010/09/10. doi: 10.1152/ajpcell.00269.2010. PubMed PMID: 20826763; PMCID: PMC2980315.

68. Gerzanich V, Woo SK, Vennekens R, Tsymbalyuk O, Ivanova S, Ivanov A, Geng Z, Chen Z, Nilius B, Flockerzi V, Freichel M, Simard JM. De novo expression of Trpm4 initiates secondary hemorrhage in spinal cord injury. Nat Med. 2009;15(2):185–91. Epub 20090125. doi: 10.1038/nm.1899. PubMed PMID: 19169264; PMCID: PMC2730968.

69. Chen X, Liu K, Lin Z, Huang K, Pan S. Knockout of Transient Receptor Potential Melastatin 4 Channel Mitigates Cerebral Edema and Neuronal Injury After Status Epilepticus in Mice. J Neuropathol Exp Neurol. 2020;79(12):1354–64. doi: 10.1093/jnen/nlaa134. PubMed PMID: 33186453.

