## Supplemental File for "IP_3_R-TRPM4 Coupling Determines the Spatial Reach of Pericyte-Mediated Capillary Constriction"

#### SUPPLEMENTAL MATERIAL

##### Model Overview

The two-projection model is designed to simplify the calcium dynamics associated with compartmentalized pericyte projections [4, 6]. With each projection operating separately, we examined how signals propagate between projections. $Ca^{2+}$ influx in Projection 1 (P1) activates TRPM4, causing membrane depolarization [7, 10, 12, 13, 15] that spreads to Projection 2 (P2) via axial resistance (1 MΩ) and activates voltage-gated $Ca^{2+}$ channels (VGCCs) [8, 9, 11], driving $Ca^{2+}$ influx into the distal projection. $Ca^{2+}$ buffering is incorporated in each projection to maintain physiologically relevant intracellular $Ca^{2+}$ concentrations [16]. All simulations were implemented in Python using forward Euler integration (dt = 1 × $10^{-4}$ s) [17, 18].

We simulated two conditions. In the first, we varied the $Ca^{2+}$ input amplitude in P1 to understand how different levels of $Ca^{2+}$ affect TRPM4 activation and downstream signaling. The $Ca^{2+}$ input to P1 is modeled using a Gaussian waveform [13, 17, 18] centered at a specified peak time (peak_time = 5 s, duration = 0.5 s), producing a transient $Ca^{2+}$ signal that mimics the kinetics of $IP_3R$-mediated $Ca^{2+}$ release [1, 2, 13]. In the second, we varied TRPM4 channel density, where $Ca^{2+}$ in P1 rises linearly from a baseline of 0.1 μM at a rate of 0.05 μM/s and is capped at 1.0 μM to maintain physiological concentrations [16].

$$calcium_{influx(P1)} = Amplitude * e^{-\frac{(time - peak)^2}{2*duration^2}}$$

Amplitude sets the peak $Ca^{2+}$ concentration of the Gaussian waveform, allowing us to simulate varied amounts of $Ca^{2+}$ release from different sources (e.g., VGCC, $IP_3R$) [1, 2, 4, 8, 13]. By varying amplitude, we can observe how TRPM4 conductance [12, 13, 15] changes in response to different $Ca^{2+}$ levels. Peak time defines when $Ca^{2+}$ concentration reaches its maximum in P1, and varying this parameter allows us to examine how the timing of $Ca^{2+}$ input affects TRPM4 activation. Duration determines how long the $Ca^{2+}$ influx is sustained in P1, allowing us to compare TRPM4 conductance under prolonged versus transient $Ca^{2+}$ signals [1, 17].

The TRPM4 conductance in P1 is determined by a bell-shaped Gaussian function [13, 15], where activation depends on the $Ca^{2+}$ concentration in P1. TRPM4 conductance reaches its peak at an optimal $Ca^{2+}$ concentration [12, 19] ($Ca^{2+}$_optimum) and diminishes as $Ca^{2+}$ deviates from this value in either direction [12, 20].

$$TRPM4_{conductance} = TRPM4(N) * Max.TRPM4\_conductance * e^{-\frac{(calcium_{influx(P1)} - calcium_{optimum})^2}{2*standard\ deviation^2}}$$

TRPM4(N) represents the number of TRPM4 channels, which scales the overall conductance and determines the total depolarizing current [12, 15].

Max.TRPM4_conductance is the maximum conductance of a single TRPM4 channel, set to 30 pS based on literature-measured values [12, 19]. $Ca^{2+}$_optimum is the $Ca^{2+}$ concentration at which TRPM4 conductance peaks, as determined from the literature [19, 20], and defines the channel's sensitivity to $Ca^{2+}$ influx. Standard deviation defines the width of the $Ca^{2+}$ activation range around this optimum [12, 13].

Membrane potential in P1 is updated at each time step using the forward Euler method [17, 18], driven by the TRPM4 current [12, 13]. The equation for membrane depolarization [21] is:

$$V_m = V_m(i-1) + \Delta t \cdot (- \frac{TRPM4_{conductance}(i) \cdot (V_m(i-1) - E_{TRPM4})}{C_m}$$

Vm is updated at each time step Δt from its previous value $Vm_{i-1,}$ driven by the TRPM4 current [18, 21]. ETRPM4 (reversal potential = 0 mV) sets the direction and magnitude of depolarization [12, 15], and Cm (membrane capacitance = 1 μF) moderates the rate of voltage change [18, 21].
As membrane depolarization spreads from P1 to P2 [4, 5], VGCCs are activated following a sigmoidal voltage dependence [5, 8, 21], with VGCC conductance increasing proportionally with depolarization [8, 9].

$$VGCC_{conductance} = \frac{VGCC_{scaling\_factor} \cdot Max.TRPM4\_conductance}{1 + e^{\frac{V_{1/2} - V_m}{k}}}$$

$VGCC_{scaling\_factor}$ adjusts the maximum VGCC conductance that the model can reach to ensure it has values closer to measured values [5, 9]. $V_{1/2}$ is the half-activation voltage of the membrane depolarization at which the VGCC conductance is at 50% of its maximum value ($V_{1/2}$) [5, 8, 21], this is essential to set the voltage sensitivity of VGCC. ‘k’ is the slope of the curve [18, 21].

$Ca^{2+}$ influx into P2 is derived from Ohm’s law, incorporating VGCC conductance and intracellular buffering, consistent with established approaches to modeling $Ca^{2+}$ dynamics [16-18]. Buffering is included to prevent nonphysiological $Ca^{2+}$ accumulation [2, 16] and to reflect the role of intracellular $Ca^{2+}$ buffering proteins in maintaining free $Ca^{2+}$ within a physiological range [1, 16].

$${Ca^{2+}}_{projection\ 2} = \frac{\Delta t \cdot (- {VGCC_{conductance}}/{C_m})}{1 + buffer_{capacity}(P2)}$$

Supplementary Figure 4 shows the experimental $Ca^{2+}$ event data that informed the choice of input waveforms for the model. Supplementary Figures 5–9 show the effect of varying $Ca^{2+}$ amplitude on TRPM4 conductance, VGCC activation, and $Ca^{2+}$ influx in P2, with 0.5 μM included as a reference condition in each figure.
Sup. Fig. 2 shows that at low-to-moderate $Ca^{2+}$ amplitudes (0.5, 0.6, and 0.7 μM), TRPM4 conductance is highest at 0.6 μM, just above the optimal $Ca^{2+}$ concentration for TRPM4 activation [12, 19]. As amplitude increases beyond this point, TRPM4 conductance decreases [13, 15], consistent with the bell-shaped $Ca^{2+}$ sensitivity of the channel [12, 13, 20]. Sup. Fig. 9 shows that at moderate-to-high $Ca^{2+}$ amplitudes (0.5, 1.0, and 1.5 μM), TRPM4 activates faster but with progressively lower integrated conductance, reducing VGCC activation and $Ca^{2+}$ influx into P2 [5, 12, 13]. Sup. Fig. 1 extends this to high amplitudes (0.5, 1.5, and 2.0 μM), where the TRPM4 conductance window narrows further and the 1.5 and 2.0 μM conditions produce nearly indistinguishable VGCC profiles in P2 [13, 15]. Sup. Fig. 2 shows that at very high amplitudes (0.5, 2.5, and 3.0 μM), only a brief low-conductance TRPM4 spike is produced at stimulus onset, with negligible VGCC activation and $Ca^{2+}$ influx [9, 12, 13] in P2. Sup. Fig. 9 shows that at extreme amplitudes (10, 15, 20, and 25 μM), $Ca^{2+}$ saturates the physiological cap instantaneously, TRPM4 conductance is abolished, and $Ca^{2+}$ influx into P2 is negligible [13, 15, 16] across all conditions.

The simulations show that moderate $Ca^{2+}$ levels (~0.5–0.6 μM) produce the most sustained TRPM4 activation [12, 19], which in turn drives efficient VGCC activation [5, 8] and $Ca^{2+}$ influx in P2 [5, 9].

At higher $Ca^{2+}$ amplitudes (above 0.6 μM), TRPM4 activates faster but with shorter duration and lower total conductance [12, 13], reducing VGCC activation and $Ca^{2+}$ influx into P2 [5, 9]. At lower $Ca^{2+}$ amplitudes (below 0.5 μM), TRPM4 is insufficiently activated [12, 19], resulting in poor VGCC activation and minimal $Ca^{2+}$ propagation to P2. These results suggest that both under- and over-activation of TRPM4 impair projection coupling [13, 15], and that the channel operates most effectively within a narrow $Ca^{2+}$ concentration range [12, 20].

Simulations with varying TRPM4 channel densities show that even a small number of functional TRPM4 channels is sufficient [12, 22] to establish electrical coupling between projections. This suggests that pericytes can regulate their contractile response by modulating TRPM4 expression [11, 12, 14, 15, 22], which may have implications for both physiological signaling and pathological conditions [4, 7, 15, 22, 23].

TRPC3 is a DAG-activated non-selective cation channel [24-26]. To examine how TRPC3 compares to TRPM4 in coupling $Ca^{2+}$ signals between projections, we implemented the same two-compartment model with TRPC3 as the depolarizing channel in P1. The core architecture, axial resistance (10 MΩ) [17, 18], and VGCC equations in P2 were kept identical to the TRPM4 model so that differences in coupling efficiency could be attributed solely to the channel properties. In the model, GPCR stimulation is represented as a rectangular pulse (t = 0 to 0.5 s) [26, 27] that drives DAG production and degradation [24, 27] according to the following kinetics:

$$\frac{dDAG}{dt} = k_{prod} * GPCR\ (t) * \left(1 - \frac{DAG}{DAG_{max}}\right) - \left(k_{deg} + \ k_{buf}\right) * DAG$$

where $k_{prod}$ is the rate of DAG production (5.0 $s^{-1}$) [26-28], $k_{deg}$ is the rate of DAG degradation (1.0 $s^{-1}$) [27, 28], $k_{buf}$ is the rate of DAG buffering coefficient (0.5 $s^{-1}$) [26, 27], and $DAG_{max}$ is the maximum DAG concentration the system can reach (2.0 μM) [24, 27]. Not all DAG is available to activate TRPC3, and a fraction is sequestered by buffering [25, 28]. The free DAG available for channel activation is calculated as:

$$DAG_{free} = DAG * \left(\frac{k_{buf}}{\left(k_{deg} + \ k_{buf}\right)}\right)$$

Free DAG then activates TRPC3 via a Hill equation:

$$P_o = \frac{DAG_{free}^2}{(K_{DAG}^2 + DAG_{free}^2)}$$

where K_DAG is the half-maximal DAG concentration for TRPC3 activation (0.9 μM), consistent with experimentally measured DAG sensitivity of TRPC3 [24, 25] and n = 2 is the Hill coefficient [24, 27, 29], reflecting the cooperative nature of DAG binding.

TRPC3 conductance is scaled by channel density and capped at 60 pS per channel [24, 25], consistent with the reported single-channel conductance range of 40–60 pS, with a reversal potential of 0 mV [24, 25]. The TRPC3 current and the axial current from P2 together drive changes in membrane potential in P1, with a membrane resistance of 5,000 MΩ [17, 18]. The membrane potential in P2 is clamped at −10 mV to prevent nonphysiological depolarization [18]. VGCC conductance in P2 is modeled using a Boltzmann activation function [8, 21], where V½ is the half-activation voltage (−45 mV) [5, 8], k is the slope factor [18, 21], and $g_{VGCCmax}$ is the maximum VGCC conductance (30 pS) [5, 9].

$Ca^{2+}$ influx into P2 is governed by:

$$\frac{dCa}{dt} = I_{Ca(P2)} - buffering_{rate} * (Ca_{P2} - Ca_{rest})$$

where $buffering_{rate}$ is the rate at which $Ca^{2+}$ is removed from P2 (0.1 $s^{-1}$) [16, 17] and $Ca_{rest}$ is the resting $Ca^{2+}$ concentration (0.05 μM) [2, 16]. Sup. Fig. 1 shows the TRPC3 activation pathway schematic and simulated TRPC3 conductance (A), VGCC conductance in P2 (B), and $Ca^{2+}$ concentration in P2 (C) across channel densities of 1, 5, 10, 15, and 50, demonstrating that meaningful VGCC activation requires substantially higher TRPC3 densities than TRPM4 [7, 12, 24, 25]. Channel density was further varied from 100 to 2000 in Sup. Fig. 2 to establish the upper boundary of TRPC3-driven VGCC activation, confirming that even at extreme densities TRPC3 is a far less efficient mediator of projection coupling than TRPM4 [7, 12, 13, 22]. The time step, initial membrane potential, and physiological $Ca^{2+}$ cap is the same as in the TRPM4 simulations.

ANO1 is a $Ca^{2+}$-activated $Cl^-$ channel [30-32]. To examine how ANO1 (TMEM16A) compares to TRPM4 and TRPC3 in coupling $Ca^{2+}$ signals between projections, we implemented the same two-compartment model with ANO1 as the depolarizing channel in P1. The core architecture, axial resistance (10 MΩ), and VGCC equations in P2 were kept identical to the TRPC3 model. Unlike the TRPC3 model where DAG drives channel activation, ANO1 is directly activated by $Ca^{2+}$ in P1 [30-32].

The $Ca^{2+}$ input to P1 is modeled as an exponential decay transient to mimic a physiological $Ca^{2+}$ event [1, 2, 17]:

$$Ca_{P1}(t) = Ca_{rest} + \left(Ca_{peak} - Ca_{rest}\right) * exp^{\left(-\frac{t}{\tau}\right)}$$

with $Ca_{peak}$ = 1.0 μM [2, 16], τ = 0.5 s [1, 17], $Ca_{rest}$ = 0.05 μM [2, 16].

ANO1 open probability:

$$P_o = \frac{Ca^n}{(Kd^n + Ca^n)}$$

where Kd is the $Ca^{2+}$ concentration at half-maximal ANO1 activation (0.8 μM), consistent with the experimentally reported values [30-32] and n is the Hill coefficient [31, 32], reflecting the cooperative nature of $Ca^{2+}$ binding to ANO1.

ANO1 single-channel conductance is set to 2 pS, consistent with the reported range of 1–3 pS [30-32]. The reversal potential for ANO1-mediated $Cl^-$ current (E_Cl) is −30 mV [30-32], which is depolarizing relative to the resting membrane potential of −70 mV [21, 31, 32].

The ANO1 current is therefore calculated [21, 31] as:

$$I_{ANO1} = g_{ANO1} * (E_{Cl} - V_{m(P1)})$$

As with the TRPC3 model, membrane potential in P2 is clamped at −10 mV, and VGCC conductance and $Ca^{2+}$ buffering in P2 are calculated using the same equations and parameter values. Channel density was varied from 50 to 10000. Across this range, ANO1-driven depolarization was not sufficient to activate VGCCs in P2 [31, 32], indicating that ANO1 is a poor mediator [12, 32, 33] of electrical coupling between projections compared to TRPM4.

The simulation function pericyte projection(amplitude) computes the following at each time step: $Ca^{2+}$ concentration in P1, TRPM4 conductance, membrane potential in P1 and P2, VGCC conductance in P2, and $Ca^{2+}$ influx into P2. The code for this function is provided below.

### SUPPLEMENTAL FIGURES

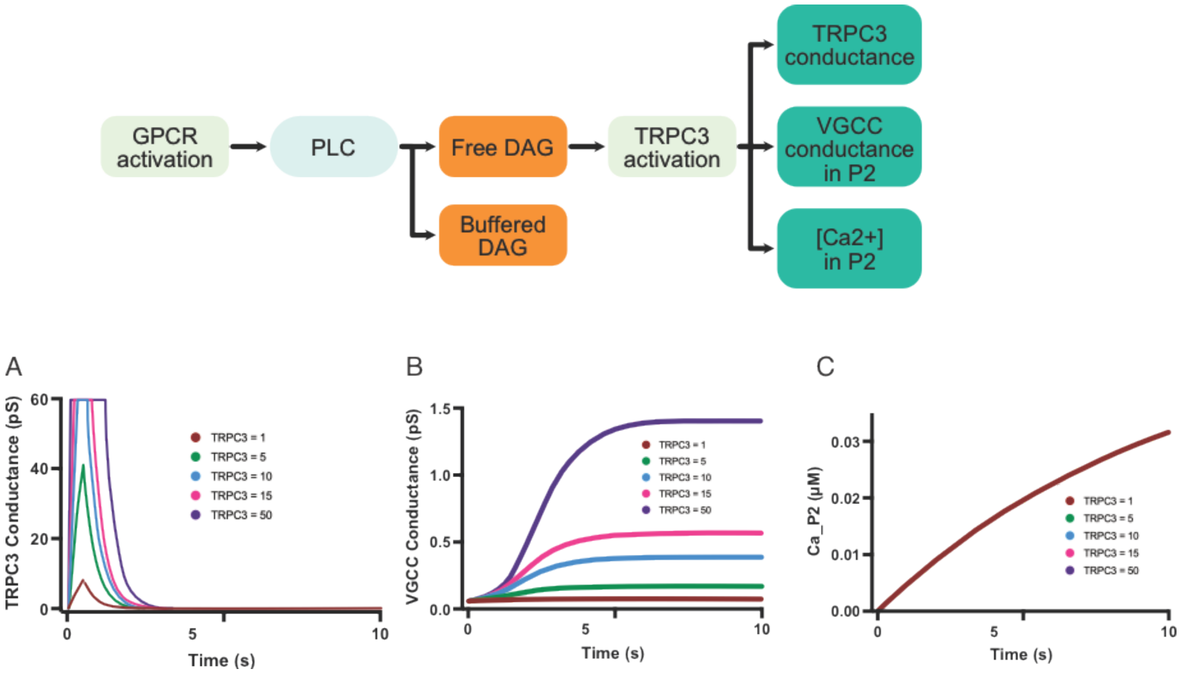


**Sup. Figure 1: TRPC3 activation pathway and downstream effects on projection coupling (Channel densities = 1, 5, 10, 15, and 50).** GPCR activation stimulates phospholipase C (PLC), leading to the generation of free diacylglycerol (DAG) and its buffered form. Free DAG activates TRPC3 channels via a Hill equation, modulating membrane potential in P1 and driving downstream signaling in P2. The schematic illustrates the full activation pathway: GPCR → PLC → free DAG → TRPC3 activation → VGCC conductance in P2 → [$Ca^{2+}$] in P2. (A) TRPC3 conductance over time for channel densities of 1, 5, 10, 15, and 50: conductance rises rapidly at stimulus onset and decays as free DAG is degraded, with higher densities producing proportionally greater conductance. (B) VGCC conductance in P2: meaningful VGCC activation is only observed at higher TRPC3 densities, in contrast to TRPM4 which achieves near-maximal VGCC coupling at low channel numbers. (C) $Ca^{2+}$ concentration in P2 over time: accumulation is density-dependent but remains low across all tested conditions. These results demonstrate that TRPC3 requires substantially higher channel densities than TRPM4 to produce comparable downstream VGCC activation and $Ca^{2+}$ influx in P2.

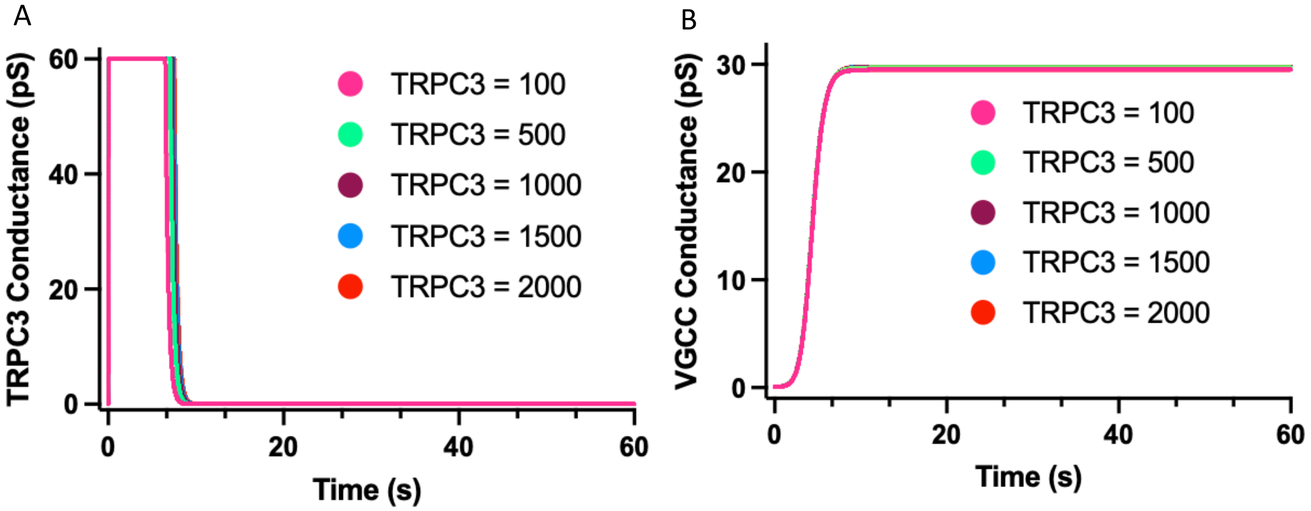


**Sup. Figure 2: High TRPC3 channel densities are required for saturation of VGCC conductance (Channel densities = 100, 500, 1000, 1500, and 2000).** Extending the TRPC3 density sweep to 100–2000 channels establish the upper boundary of TRPC3-driven VGCC activation. All model parameters are identical to Sup. Fig. 1. (A) TRPC3 conductance over time: at densities of 100 and above, total TRPC3 conductance rapidly saturates the 60 pS cap during the 0–0.5 s DAG stimulus window, with all conditions producing nearly indistinguishable conductance profiles. (B) VGCC conductance in P2: even at these extreme densities, meaningful VGCC activation is delayed and converges at saturation only above 500 channels. Taken together with Sup. Fig. 1, these results confirm that TRPC3 requires channel densities orders of magnitude greater than TRPM4 to achieve equivalent VGCC activation in P2.

**Sup. Figure 3: High ANO1 channel densities are required for saturation of VGCC conductance (Channel densities = 50, 100, 500, 1000, 5000, and 10000).** Predicted VGCC conductance in the distal pericyte projection (P2) over time for increasing ANO1 channel densities (50, orange; 100, red; 500, green; 1000, purple; 5000, cyan; 10000, pink). All model parameters are identical to those described in Fig. 2A. At low-to-moderate densities (50–100 channels), ANO1 fails to produce appreciable VGCC conductance. At 500–1000 channels, a gradual and incomplete rise is observed, failing to reach saturation within the 60-second simulation window. Only at extreme densities (5000–10000 channels) does ANO1-driven VGCC conductance approach saturation (~30 pS). These results demonstrate that ANO1 requires channel densities orders of magnitude greater than TRPM4 to achieve equivalent VGCC activation in P2, making it an unlikely primary mediator of inter-projection signal propagation under physiological conditions.

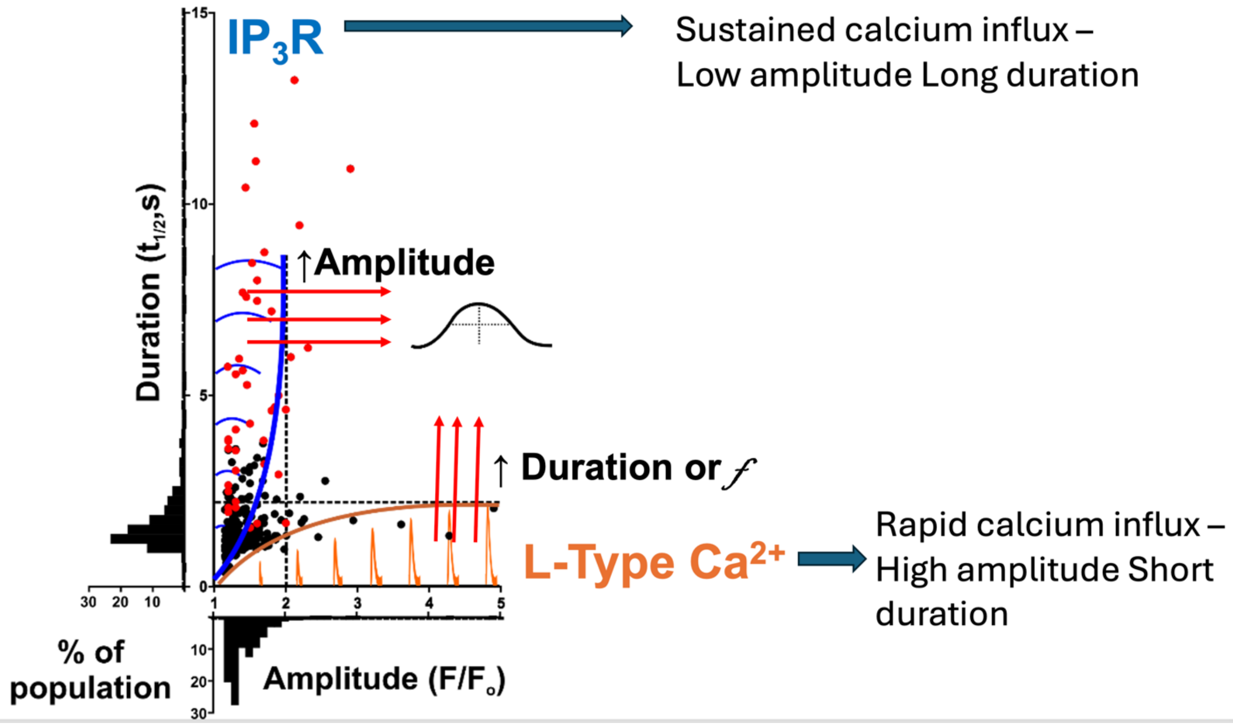


**Sup. Figure 4: Distinct $Ca^{2+}$ events mediated by $IP_3R$ and VGCCs.** Scatter plot of $Ca^{2+}$ events illustrating the relationship between amplitude ($\Delta F/F_0$) and duration ($t_{1/2}$) of individual calcium transients. Events are categorized based on their distinct profiles: (1) $IP_3R$-mediated $Ca^{2+}$ events exhibit sustained $Ca^{2+}$ influx with low amplitude and long duration (blue arrows and curve), and (2) L-type $Ca^{2+}$ channel-mediated calcium events display rapid calcium influx with high amplitude and short duration (orange arrows and curve). Insets represent schematic traces highlighting the differences in amplitude and duration between the two event types. Histograms along the axes display the frequency distributions of calcium event amplitudes and durations. These results illustrate the distinct kinetics and populations of calcium signaling pathways mediated by $IP_3Rs$ and L-type $Ca^{2+}$ channels.

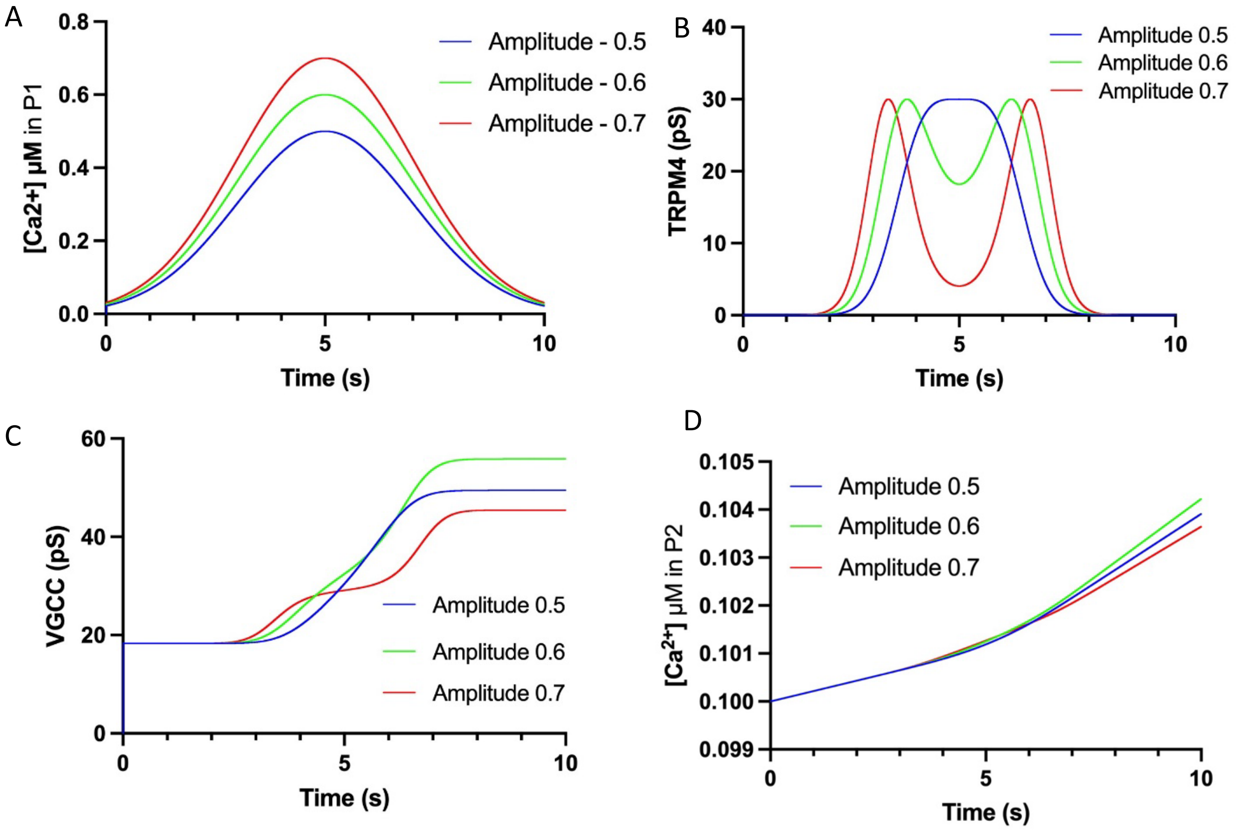


**Sup. Figure 5**: **Effects of low-to-moderate $Ca^{2+}$ amplitudes on TRPM4 conductance (Amplitudes = 0.5, 0.6, and 0.7 µM).** This figure illustrates how varying $Ca^{2+}$ concentration in Projection P1 affects TRPM4 conductance and downstream signaling. **(A)** $Ca^{2+}$ concentration in P1: higher amplitudes result in greater $Ca^{2+}$ influx. **(B)** TRPM4 conductance: since the optimal activation range for TRPM4 is ~0.5 µM $Ca^{2+}$, the conductance curve has a single peak. The maximum conductance is observed at 0.6 µM, as measured by the area under the conductance curve, indicating the optimal range for TRPM4 activation. Increasing amplitude beyond this point decreases TRPM4 activation. (C) VGCC conductance in P2: maximum VGCC conductance is also observed at 0.6 µM, consistent with the TRPM4 conductance profile. (D) $Ca^{2+}$ influx into P2 is greatest at 0.6 µM, reflecting the downstream effect of optimal TRPM4 activation.

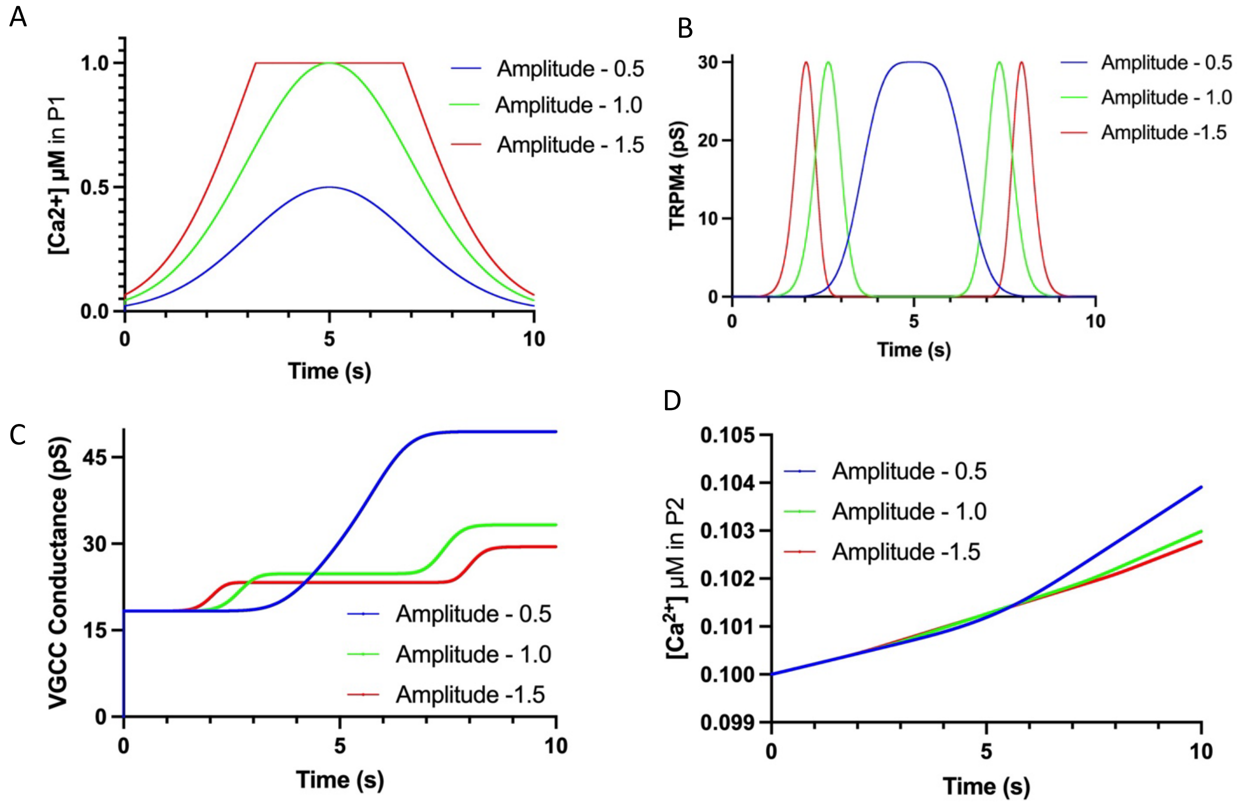


**Sup. Figure 6: Effects of moderate-to-high $Ca^{2+}$ amplitudes on TRPM4 activation and conductance (Amplitudes = 0.5, 1.0, and 1.5 µM).** Higher amplitudes result in increased $Ca^{2+}$ concentration in P1, however sustained elevated amplitude results in less TRPM4 activation. (A) $Ca^{2+}$ concentration in P1: higher amplitudes produce faster-rising and higher-peak $Ca^{2+}$ transients. (B) TRPM4 conductance: the area under the conductance curve reduces significantly at higher amplitudes — TRPM4 channels activate more quickly but with much lower total conductance, as the $Ca^{2+}$ signal moves away from the optimal activation range. (C) VGCC conductance in P2: VGCC conductance is highest at the 0.5 µM reference condition and decreases progressively with increasing amplitude. (D) $Ca^{2+}$ influx into P2 follows the same trend, with decreasing accumulation at higher amplitudes. These results illustrate how supra-optimal $Ca^{2+}$ levels reduce TRPM4 activation and downstream VGCC conductance, resulting in decreased $Ca^{2+}$ influx into Projection P2.

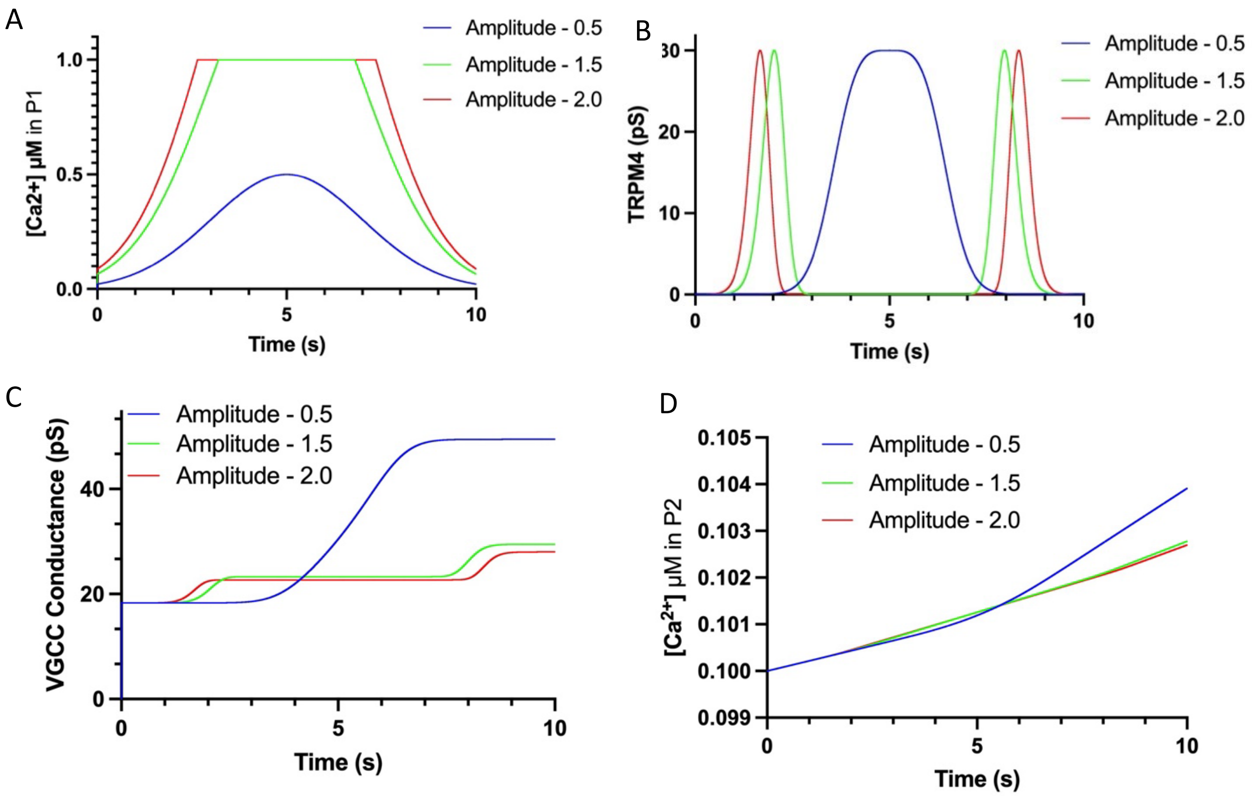


**Sup. Figure 7: Effects of high $Ca^{2+}$ amplitudes on TRPM4 activation and conductance (Amplitudes = 0.5, 1.5, and 2.0 μM).** High amplitudes result in faster activation of TRPM4 channels but lower total TRPM4 conductance. (A) $Ca^{2+}$ concentration in P1: at 1.5 and 2.0 μM, $Ca^{2+}$ rapidly saturates the physiological cap. (B) TRPM4 conductance: the area under the conductance curve decreases significantly at higher amplitudes. (C) VGCC conductance in P2: the 1.5 and 2.0 μM conditions produce nearly indistinguishable VGCC profiles, both substantially lower than the 0.5 μM reference. (D) $Ca^{2+}$ influx into P2 is reduced accordingly. These results show that high $Ca^{2+}$ amplitudes are less effective at sustaining TRPM4 activation and downstream VGCC coupling than the optimal amplitude.

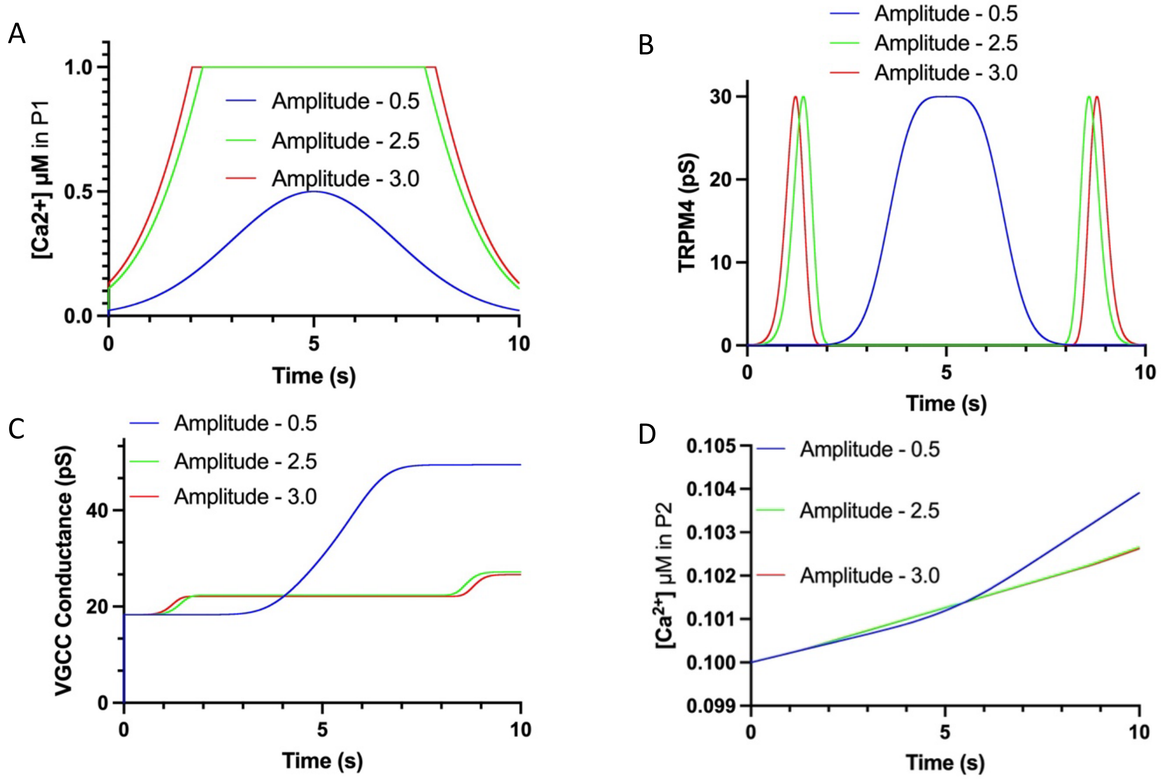


**Sup. Figure 8: Effects of very high $Ca^{2+}$ amplitudes on TRPM4 activation and conductance (Amplitudes = 0.5, 2.5, and 3.0 µM).** With very high amplitudes, $Ca^{2+}$ peak is achieved quickly, however TRPM4 activation is quicker but with much lower conductance. (A) $Ca^{2+}$ concentration in P1: at 2.5 and 3.0 µM, $Ca^{2+}$ saturates the physiological cap almost instantaneously. (B) TRPM4 conductance: only a brief, low-conductance spike is produced at stimulus onset; the total area under the conductance curve is markedly reduced compared to the 0.5 µM reference. (C) VGCC conductance in P2: increasing amplitudes beyond the optimum do not produce meaningful increases in VGCC activation; the 2.5 and 3.0 µM conditions are nearly indistinguishable. (D) $Ca^{2+}$ influx into P2 is negligible for both high-amplitude conditions and shows no further increase with amplitude. These results show that the effect of amplitude on TRPM4 activation and downstream coupling diminishes after an optimal level.

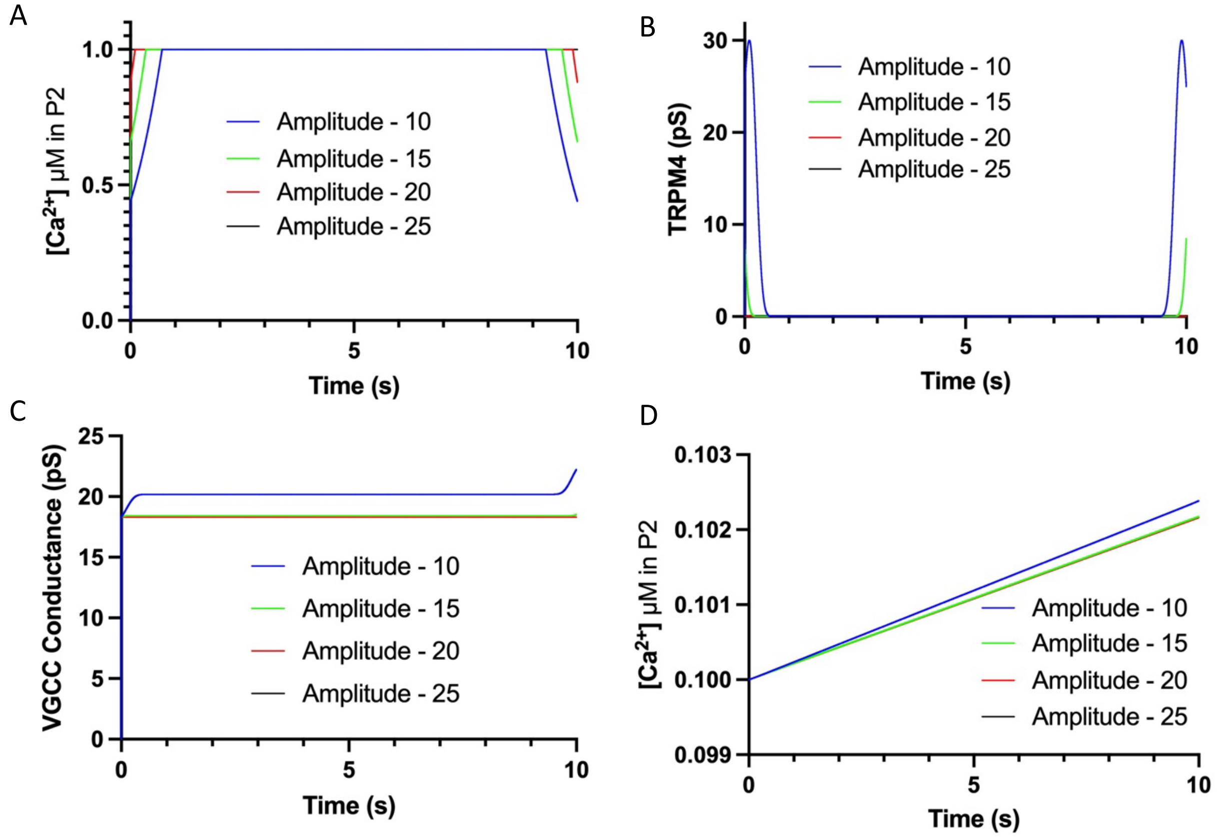


**Sup. Figure 9: Effects of extreme $Ca^{2+}$ amplitudes on TRPM4 activation and conductance (Amplitudes = 10, 15, 20, and 25 µM).** At extreme amplitudes, $Ca^{2+}$ reaches the physiological cap instantaneously at the start of the simulation. (A) $Ca^{2+}$ concentration in P1: all four conditions saturate the 1.0 µM physiological cap immediately, with no transient kinetics. (B) TRPM4 conductance: conductance is effectively abolished across all extreme amplitudes, as the $Ca^{2+}$ signal dwells entirely outside the optimal activation range for the duration of the simulation. (C) VGCC conductance in P2: VGCC activation remains low and uniform across all four conditions, with no amplitude-dependent differences. (D) $Ca^{2+}$ influx into P2 is negligible across all conditions. These results define the upper boundary of productive TRPM4-mediated signaling — beyond extreme $Ca^{2+}$ levels, increasing amplitude provides no benefit and eliminates projection coupling entirely.

#### REFERENCES FOR SUPPLEMENTAL


1 Berridge, M.J.: ‘Inositol trisphosphate and calcium signalling’, Nature, 1993, 361, (6410), pp. 315-325
2 Berridge, M.J., Bootman, M.D., and Roderick, H.L.: ‘Calcium signalling: dynamics, homeostasis and remodelling’, Nat Rev Mol Cell Biol, 2003, 4, (7), pp. 517-529
3 Earley, S., and Brayden, J.E.: ‘Transient receptor potential channels in the vasculature’, Physiol Rev, 2015, 95, (2), pp. 645-690
4 Hartmann, D.A., Berthiaume, A.A., Grant, R.I., Harrill, S.A., Koski, T., Tieu, T., McDowell, K.P., Faino, A.V., Kelly, A.L., and Shih, A.Y.: ‘Brain capillary pericytes exert a substantial but slow influence on blood flow’, Nat Neurosci, 2021, 24, (5), pp. 633-645
5 Klug, N.R., Sancho, M., Gonzales, A.L., Heppner, T.J., O'Brien, R.I.C., Hill-Eubanks, D., and Nelson, M.T.: ‘Intraluminal pressure elevates intracellular calcium and contracts CNS pericytes: Role of voltage-dependent calcium channels’, Proc Natl Acad Sci U S A, 2023, 120, (9), pp. e2216421120
6 Hall, C.N., Reynell, C., Gesslein, B., Hamilton, N.B., Mishra, A., Sutherland, B.A., O'Farrell, F.M., Buchan, A.M., Lauritzen, M., and Attwell, D.: ‘Capillary pericytes regulate cerebral blood flow in health and disease’, Nature, 2014, 508, (7494), pp. 55-60
7 Gonzales, A.L., Klug, N.R., Moshkforoush, A., Lee, J.C., Lee, F.K., Shui, B., Tsoukias, N.M., Kotlikoff, M.I., Hill-Eubanks, D., and Nelson, M.T.: ‘Contractile pericytes determine the direction of blood flow at capillary junctions’, Proc Natl Acad Sci U S A, 2020, 117, (43), pp. 27022-27033
8 Catterall, W.A.: ‘Voltage-gated calcium channels’, Cold Spring Harb Perspect Biol, 2011, 3, (8), pp. a003947
9 Knot, H.J., and Nelson, M.T.: ‘Regulation of arterial diameter and wall [Ca2+] in cerebral arteries of rat by membrane potential and intravascular pressure’, J Physiol, 1998, 508 ( Pt 1), (Pt 1), pp. 199-209
10 Gonzales, A.L., and Earley, S.: ‘Endogenous cytosolic Ca(2+) buffering is necessary for TRPM4 activity in cerebral artery smooth muscle cells’, Cell Calcium, 2012, 51, (1), pp. 82-93
11 Gonzales, A.L., and Earley, S.: ‘Regulation of cerebral artery smooth muscle membrane potential by Ca(2)(+)-activated cation channels’, Microcirculation, 2013, 20, (4), pp. 337-347
12 Launay, P., Fleig, A., Perraud, A.L., Scharenberg, A.M., Penner, R., and Kinet, J.P.: ‘TRPM4 is a Ca2+-activated nonselective cation channel mediating cell membrane depolarization’, Cell, 2002, 109, (3), pp. 397-407
13 Gaur, N., Hof, T., Haissaguerre, M., and Vigmond, E.J.: ‘Propagation Failure by TRPM4 Overexpression’, Biophys J, 2019, 116, (3), pp. 469-476
14 Mughal, A., Harraz, O.F., Gonzales, A.L., Hill-Eubanks, D., and Nelson, M.T.: ‘PIP(2) Improves Cerebral Blood Flow in a Mouse Model of Alzheimer's Disease’, Function (Oxf), 2021, 2, (2), pp. zqab010
15 Abriel, H., Syam, N., Sottas, V., Amarouch, M.Y., and Rougier, J.S.: ‘TRPM4 channels in the cardiovascular system: physiology, pathophysiology, and pharmacology’, Biochem Pharmacol, 2012, 84, (7), pp. 873-881

16 Neher, E., and Augustine, G.J.: ‘Calcium gradients and buffers in bovine chromaffin cells’, J Physiol, 1992, 450, pp. 273-301
17 Schuster, S., Marhl, M., and Hofer, T.: ‘Modelling of simple and complex calcium oscillations. From single-cell responses to intercellular signalling’, Eur J Biochem, 2002, 269, (5), pp. 1333-1355
18 Destexhe, A., and Huguenard, J.R.: ‘Nonlinear thermodynamic models of voltage-dependent currents’, J Comput Neurosci, 2000, 9, (3), pp. 259-270
19 Vennekens, R., and Nilius, B.: ‘Insights into TRPM4 function, regulation and physiological role’, Handb Exp Pharmacol, 2007, (179), pp. 269-285
20 Uchida, K.: ‘TRPM3, TRPM4, and TRPM5 as thermo-sensitive channels’, J Physiol Sci, 2024, 74, (1), pp. 43
21 Hodgkin, A.L., and Katz, B.: ‘The effect of sodium ions on the electrical activity of giant axon of the squid’, J Physiol, 1949, 108, (1), pp. 37-77
22 Hariharan, A., Weir, N., Robertson, C., He, L., Betsholtz, C., and Longden, T.A.: ‘The Ion Channel and GPCR Toolkit of Brain Capillary Pericytes’, Front Cell Neurosci, 2020, 14, pp. 601324
23 Eltanahy, A.M., Aupetit, A., Buhr, E.D., Van Gelder, R.N., and Gonzales, A.L.: ‘Light-sensitive Ca(2+) signaling in the mammalian choroid’, Proc Natl Acad Sci U S A, 2024, 121, (46), pp. e2418429121
24 Hofmann, T., Obukhov, A.G., Schaefer, M., Harteneck, C., Gudermann, T., and Schultz, G.: ‘Direct activation of human TRPC6 and TRPC3 channels by diacylglycerol’, Nature, 1999, 397, (6716), pp. 259-263
25 Venkatachalam, K., Zheng, F., and Gill, D.L.: ‘Regulation of canonical transient receptor potential (TRPC) channel function by diacylglycerol and protein kinase C’, J Biol Chem, 2003, 278, (31), pp. 29031-29040
26 Berridge, M.J.: ‘Inositol trisphosphate and calcium signalling mechanisms’, Biochim Biophys Acta, 2009, 1793, (6), pp. 933-940
27 Gonzales, D.T., Schuhmacher, M., Lennartz, H.M., Iglesias-Artola, J.M., Kuhn, S.M., Barahtjan, P., Zechner, C., and Nadler, A.: ‘Quantifying single-cell diacylglycerol signaling kinetics after uncaging’, Biophys J, 2024, 123, (7), pp. 921-927
28 Rhee, S.G.: ‘Regulation of phosphoinositide-specific phospholipase C’, Annu Rev Biochem, 2001, 70, pp. 281-312
29 Mederos, Y.S.M., Storch, U., and Gudermann, T.: ‘Mechanosensitive G(q/11) Protein-Coupled Receptors Mediate Myogenic Vasoconstriction’, Microcirculation, 2016, 23, (8), pp. 621-625
30 Caputo, A., Caci, E., Ferrera, L., Pedemonte, N., Barsanti, C., Sondo, E., Pfeffer, U., Ravazzolo, R., Zegarra-Moran, O., and Galietta, L.J.: ‘TMEM16A, a membrane protein associated with calcium-dependent chloride channel activity’, Science, 2008, 322, (5901), pp. 590-594
31 Hartzell, H.C., and Whitlock, J.M.: ‘TMEM16 chloride channels are two-faced’, J Gen Physiol, 2016, 148, (5), pp. 367-373
32 Korte, N., Ilkan, Z., Pearson, C.L., Pfeiffer, T., Singhal, P., Rock, J.R., Sethi, H., Gill, D., Attwell, D., and Tammaro, P.: ‘The Ca2+-gated channel TMEM16A amplifies capillary pericyte contraction and reduces cerebral blood flow after ischemia’, J Clin Invest, 2022, 132, (9)

33 Al-Hosni, R., Kaye, R., Choi, C.S., and Tammaro, P.: ‘The TMEM16A channel as a potential therapeutic target in vascular disease’, Curr Opin Nephrol Hypertens, 2024, 33, (2), pp. 161-169
